# Inhibition of serine racemase prevents diabetic retinopathy

**DOI:** 10.1101/2024.06.30.601441

**Authors:** Haiyan Jiang, Piansi Zhou, Xue Jiang, Xiong Wu, Yandie Mao, Shuyi Liu, Shiqi Tang, Jing Zhou, Zhiwen Zhang, Bingbing Ren, Ge Shan, Jia Qu, Shengzhou Wu

**Author notes:** To whom correspondence should be addressed: Prof. Shengzhou Wu, Ph.D, M.D. equally contributed to the work (Haiyan Jiang, Piansi Zhou, Xue Jiang).

## Abstract

Retinal diabetic neuropathy (RDN) precedes retinal microvascular pathology in diabetic retinopathy (DR), but the therapy by antagonizing RDN to stall DR is rare. Emerging evidence including ours suggest that serine racemase (SRR) drives DR at least by promoting RDN. Thus, we explore the therapy by antagonizing SRR to stall and treat DR. We herein indicate that inhibition of SRR by oral gavage of l-aspartic acid β-hydroxymate (L-ABH), a competitive inhibitor of SRR, mitigated photoreceptor dysfunction, loss of retinal ganglion cells (RGC) and retinal endothelial cell and pericyte in db/db mice, a type II diabetes model. To dissect the mechanism, intravitreal injection of L-ABH mitigates glutamate-induced neurotoxicity in the retina. In the whole body, gastrointestinal-based delivery of L-ABH maintained euglycemia and improved glucose tolerance either in db/db mice or in diet-induced obesity mice. In conclusion, inhibition of SRR prevented retinal neurovascular abnormalities in diabetic animals through executing neuroprotection in the retina and maintaining glucose homeostasis in the system. Our study reveals a novel strategy to prevent DR.

## Introduction

Diabetic retinopathy (DR) is a vision-threatening complication of diabetes mellitus (DM), leading to retinal neurovascular pathologies and resulting in blindness in large numbers of populations worldwide [1]. Almost all type I (T1DM) and >60% type II DM (T2DM) patients develop retinopathy after twenty-year course of diabetes [2]. DR is characterized with manifold pathologies in the retinal microvasculature and staged into mild, moderate, and severe non-proliferative DR (NPDR) according to the extent of microaneurysms, retinal hemorrhages, exudates, and venous beadings while the proliferative DR (PDR) manifests retinal neovascularization, vitreous hemorrhage, fibrovascular membrane formation, and tractional retinal detachment [3]. Before overt microvascular abnormalities, DR frequently accompanies with structural deficits in the neuroretina such as apoptosis in the inner retina, reduction of thickness of inner plexiform layer (IPL) and inner nuclear layer (INL), and loss of retinal ganglion cell (RGC), cholinergic and dopaminergic amacrine cells, and gliosis [4–6]. In addition, functional deficits in DR retina are common including abnormal responses in electroretinogram (ERG) recordings, contrast sensitivity, and dark adaption [7–12]. The retinal neurodegeneration is originally assumed as a secondary sequela of retinal microvascular damage, but ample evidence report that the retinal diabetic neuropathy (RDN) occurs in type I and type II DM patients with minimal, or even without DR [13–16]. In a four-year longitudinal study, examinations with optical coherence tomography identify progressive loss of retinal nerve fiber layer (RNFL), RGC and IPL even though these patients do not manifest DR clinically [17]. Thus, retinal neurodegeneration precedes retinal microvascular abnormalities in DR.

Serine racemase (SRR) is a divalent cation-dependent allosteric enzyme with pyridoxal 5’-phosphate as an essential cofactor, synthesizing D-serine using l-serine as a substrate [18, 19]. Appropriate activity of N-methyl-D- aspartate receptor (NMDAR) depends on balanced levels of extracellular glutamate binding to GluN2 subunit and D-serine or glycine ligating to GluN1 subunit; depletion of D-serine largely mitigates NMDAR-mediated neuro- transmission and excitotoxicity [20–23] . NMDAR-mediated excitotoxicity has been suggested to significantly contribute to retinal neurodegeneration in DR. In early diabetes, glutamate transporter on Müller glia is significantly reduced by oxidative stress, disrupting glutamate homeostasis and thus increasing glutamate in diabetic retina [24, 25]. By chronic delivery of memantine, an uncompetitive antagonist of NMDAR used for treating Alzheimer’s disease, the retinal function, RGC loss, and the altered permeability of blood-retinal barrier in diabetic rats are significantly mitigated [26]. Thus, blockade of NMDAR- mediated retinal neurodegeneration may stall DR. However, most NMDAR antagonists cause unacceptable clinical side effects and fail in advanced clinical trials [27, 28]. Thus, we have explored the role of SRR in DR in order to develop therapy, aiming to block excitotoxicity with a clinically tolerated manner, with which to prevent and treat DR. In the past decade, we have indicated the promoting role of SRR in DR by demonstrating: (1) SRR is increased in the retina of streptozotocin-induced diabetic rats [29]; (2) D-serine is increased in the aqueous humor of streptozotocin-induced diabetic rats and also increased in the aqueous and vitreous humor of NPDR and PDR patients [29, 30]; (3) Loss of SRR due to mRNA decay significantly reduces RGC degeneration in mice induced by intravitreal injection of NMDA, and through the same mechanism, mitigates RGC loss in type I diabetic mice, Ins2^Akita^ mice [31, 32]; (4) D-amino acid oxidase (DAAOx), the enzyme degrading D-serine, is reduced in the retina of streptozotocin-induced diabetic rats and overexpression of DAAOx in the retina significantly prevents RGC loss and improves blood-retinal barrier in diabetic rats [33]. In this study, we found that oral gavage of l-aspartic acid β-hydroxymate (L-ABH), a competitive inhibitor of SRR, significantly improved retinal function evaluated by electroretinogram, mitigated the loss of RGC, retinal endothelial cell, and pericyte in db/db mice. In the retina, we further indicated that intravitreal injection of L-ABH significantly decreased glutamate-induced neurotoxicity. In the whole body, oral gavage of L-ABH maintained euglycemia and improved glucose intolerance.

## Materials

The following materials and antibodies were used for this study: antibodies:SRR(cat#612052,RRID:AB_399439,BD Biosciencs). α-Tubulin (cat#2144s, RRID:AB_2210548)(Cell Signaling). Brain specific homeobox/POU domain protein(Brn3a, RRID: AB_2167511), PEPCK(cat#sc-271029, RRID:AB_10610383 (Santa Cruz Biotechnology). G6Pase antibody(cat#bs-22837R, Bioss Inc). Collagenase P(cat# 11249002001), OptiPrep™ density gradient medium(cat# D1556), D-glucose(cat#G7021), L-ABH(cat# A6508)(Sigma). TNF-α(cat#C600052), IFN-γ(cat# C600059), IL-1β(cat#C600124), metformin(cat#A506198)(Sangon Biotech, Shanghai). Hoechst 33342(cat#C1028), insulin(cat# P3375), penicillin and streptomycin solution(cat#C0222)(Beyotime Biotechnology,Beijing). Krebs-ringer solution(cat#PB180347), HBSS(cat#PB180321)(Procell, Wuhan, China). KRBH buffer (cat#PH1832-D, Phygene, Fujian, China). FBS(cat#16000-044), MEM(cat# 11090081)(Gibco). Glycated hemoglobin A1c ELISA kit(cat#YS02875B)(YaJi Biological, Shanghai). Anti-mouse insulin ELISA kit (cat#SEKMO141), glycogen assay kit(cat#BC0345), the glucose assay kit(cat#BC2505)(Solarbio,Beijing). Sodium pyruvate(cat#P2256-5G) (Sigma). Tropicamide/ phenylephrine eye drops (Santen Pharmaceutical Co. Ltd).

## Research design and methods Ethics

This study followed COPE guidelines. The mice were fed *ad libitum* in a temperature and humidity-controlled and pathogen-free animal facility with a 12-h on/off cycle of automatic illumination in Wenzhou Medical University. All procedures were approved by the Institutional Review Board of Wenzhou Medical University, China (approval number# wydw2021-0085).

## Reverse-phase HPLC detection of l-/D-serine

Detection of l-/D-serine by reverse-phase HPLC was performed using methods similar to our previously established protocol [29]. The collected blood samples or aqueous humor samples were subjected to centrifugation at 12,000 x g for 10 min. The supernatants were collected and precipitated by adding equal volume of acetonitrile (cat#1.00030.4000, Merck) and strained with syringe-driven filter (cat#HP141,Sangon Biotech, Shanghai). The samples were then derivatized with a 3:7 mixture of solution A consisting of 30 mg/ml t-BOC-L-cysteine (cat#15411, Sigma) and 30 mg/ml o-phthaldialdehyde (cat# P0657, Sigma) in methanol and solution B (100 mM sodium tetraborate solution, pH 9.4). A ZORBAX Eclipse AAA column (3.5 μm, 150 x 4.6 mm, Agilent, USA) was used to separate amino acids. A linear gradient was established from 100% buffer A consisting of 0.1 M sodium acetate buffer (cat#S8750, Sigma), 7% acetonitrile, and 3% tetrahydrofuran (cat#401757, Sigma) to 100% buffer B consisting of 0.1 M sodium acetate buffer, 47% acetonitrile, and 3% tetrahydrofuran over 60 min at 0.8 ml/min. Fluorescence was monitored with 344 nM excitation and 443 nM emission. In addition to their consistent retention times, D-serine peaks were confirmed by sensitivity to D-amino acid oxidase (DAAOx) digestion.

## Electroretinogram recording

A RETI-Port 21 system with a custom-built Ganzfeld dome (Roland Consult, Wiesbaden, Germany) was used for electroretinographic recording as our previously established protocol [34]. The db/db mice were adapted to dark O/N before scotopic recording. On the day before recording, the mice were anesthetized with intraperitoneal injection of pentobarbital sodium/xylazine and the pupils were dilated with tropicamide/ phenylephrine eye drops. The mice were placed on a pre-heated platform with a positive loop electrode gently touching cornea, a negative electrode subcutaneously placing between the ears, and a reference electrode subcutaneously inserted into animal’s rump.

Scotopic ERGs were recorded at 0.01, 3,10 cd-s/m^2^ stimulus intensity with 30-s interval and five ERG recording were averaged for scotopic ERGs. Before photopic recording, the mice were exposed to steady illumination at 30 cd-s/m2 for 10 min. Photopic ERGs were recorded at 3 cd-s/m2 stimulus intensity with 0.4-s interval and fifty scan were averaged for photopic ERGs. Under photopic adaption, ERGs were also recorded under 30Hz flick light. The ERG recordings were conducted under a blinded mode.

## Spectral Domain Optical Coherence Tomography (SD-OCT) Imaging

A Micron IV SD-OCT system (Pheonix) was used for imaging. Before imaging, the mice were anesthetized with intraperitoneal injection of pentobarbital sodium/xylazine and the pupils were dilated with tropicamide/ phenylephrine eye drops. The mouse eyes were evenly illuminated and the camera was well focused on the optic disk to ensure high-quality images.

Under the mode of linear scan, the optic disk was positioned in the center of scan box and the images were automatically acquired from nasal to temporal retina. Under the mode of circular scan, the optic disk was wrapped by scan box and the retina surrounding optic disk was automatically scanned and the acquired images were saved. The images were processed by a built-in software, *insight*. The SD-OCT analysis was conducted under a blinded mode.

## Retinal flatmount and immunofluoresecence

Briefly, db/db mice were killed and the eyes were immediately enucleated and fixed in 4% paraformaldehyde. Each retina was cut with four radial incision and fixed again. After fixation, the retina was flattened and mounted on a glass slide with a coverslip to prepare as a retinal whole flatmount. The retina flatmount was blocked with 10% goat sera and further incubated with a RGC-specific transcriptional factor (Brn3a) antibody (1:200), in PBS containing 0.1% Triton X-100 at 4°C overnight. After washed, the retinal flat-mount was incubated with a 594-conjugated secondary antibody (1:500, Invitrogen) for 1 h and subject to analysis under fluorescent microscopy (Zeiss AX10). The fluorescently labeled RGCs were captured and counted separately for posterior, middle, peripheral retina with ImageJ software (NIH, Bethesda, Maryland, USA). The center, intermediate, or peripheral retina was demarcated as our previously established protocol (0.15 mm^2^) [31]. The counting for each retinal segment was averaged from 16-20 boxes and the counting was conducted under a blinded mode.

## Retinal vasculature

Periodic acid-Schiff (PAS) staining and haematoxylin counterstaining were used to indicate retinal vasculature as our previously established protocol [33]. The enucleated eyeballs were fixed in 4% paraformaldehyde for 48 h and the retina was gently peeled off and subjected to digestion in 3% trypsin dissolved in 0.2 M Tris buffer (pH 7.4) at 37°C for 1 h. The retinal vessels were separated from the neural tissue and limiting membrane, transferred to slides and dried at room temperature for PAS staining. The dried retinae were fixed in Carnoy’s solution, washed and further dipped in 0.5% periodic acid solution for 10 min. The slides were counterstained with Schiff solution. After washing, the slides were subjected to discoloration and subsequent dehydration. Finally, the slides were sealed with neutral balsam and subjected to haematoxylin staining.

## Intravitreal injection

C57BL/6J mice at the age of two months were randomly assigned to injection of saline (1 μL), glutamate (100 mM, 1 μL), and glutamate (100 mM) plus L-ABH (1mg/mL) (1 μL). Following our previously established protocol [33], after the pupil was dilated with two drops of tropicamide/phenylephrine hydrochloride, the mice were anaesthetized with intramuscular injection of 0.08 mL/10g body weight of ketamine/xylazine mixture (10 mg/kg). Under an eye surgery microscope, a Hamilton needle was used to perforate in the sclera at the superior temporal quadrant,1 mm posterior to limbus. After withdrawing the needle, a microsyringe was used to gradually inject the solution through the perforation. After injection, a sterilized cotton swab was used to press the injection site for 1 min and erythromycin eye ointment was applied to the eyelid. The mice were killed at 24 h after injection and the eyes were used to prepare retinal flatmount as the above procedure.

## OGTT, ITT, tissue isoltion, and blood sample collection from db/db mice

Male db/db homozygotes and male WT mice with BKS genetic background were purchased from GemPharmatech (JiangSu,China). The male mice with similar weights were randomly assigned to different groups. Before feeding chemicals, insulin sensitivity test (ITT) and oral glucose tolerance test (OGTT) were documented for WT and db/db mice. At the age of 12 weeks, Db/db male mice with similar weights (∼47g) were randomly divided into three groups which were fed with water, L-ABH (20 mg/kg/d), metformin (250 mg/kg/d) via oral gavage. The feeding was continued for 18 weeks until the age of 30 weeks and BKS WT mice were fed with water as a comparison. The body weight and blood glucose were documented once every other week. At the end of feeding, the mice were fasted for 14 h and D-glucose (2 g/kg) were fed via oral gavage. Blood samples were collected from tail vein, immediately before or at 15, 30, 60, and 90 min after D-glucose feeding. The blood glucose levels were measured with glucometer. After OGTT, the mice were continuously given water, L-ABH, and metformin, respectively for 1 week for the following ITT test. Before ITT test, the mice were fasted for 6 h and insulin (0.75 U/kg) were intraperitoneally injected. The blood samples were collected immediately before or at 15, 30, 60, and 90 min after injection and blood glucose levels were measured with glucometer. After one week with continuous feeding, the mice were anaesthetized with intraperitoneal injection of sodium pentobarbital (0.36 mg/10 g body weight) and xylazine (0.054 mg/10 g body weight) and the livers were extracted to examine *pepck* and *g6pase* mRNA and protein levels. Under anesthetization, blood samples were collected from eye socket for measuring HA1C and insulin. The mice were immediately killed by detachment of cervical spine. HOMA-IR was calculated with the formula: fasting glucose (mmol/L) x fasting insulin (mIU/L))/22.5.

## ITT, OGTT, tissue isolation from diet-induced obesity mice

*Srr*^-/-^ mice were generated with the CRSIPR/Cas9 technology in Biocytogen, Inc (Beijing, China) and the model was used in our very recent study[35]. *Srr*^-/-^ or littermate WT male mice with C57BL/6 background were fed high fat diet(HFD) or normal chow diet(CD) since the age of 3 weeks. The male mice with similar weights (∼25g) were randomly assigned to different groups. At the age of 15 weeks, ITT was immediately conducted after the mice were fasted for 6 h. After fasting, insulin (0.75 U/kg) was intraperitoneally injected and blood samples were collected from tail vein for measuring glucose concentration, immediately before or at 15, 30, 60, and 90 min after injection. At the age of 19 weeks, oral glucose tolerance test (OGTT) was immediately performed after the mice were fasted for 14 h. After fasting, D-glucose solution(2g/kg) was fed via oral gavage and blood samples were collected for glucose examination, immediately before or at 30, 60, 90, and 120 min after glucose feeding.

To test the effect of L-ABH on DIO mice, C57BL/6 male mice were fed HFD. At the third day of HFD feeding, the mice were randomly assigned to two groups with one group fed with water and the other with L-ABH(20 mg/kg/d).

The body weight and random blood glucose were documented once a week. At the age of 15 weeks, ITT was conducted as aforementioned. At the age of 19 weeks, OGTT was conducted as above. After the operation, the mice were immediately killed by decapitation.

## Primary hepatocyte culture

Adult WT and *Srr*^-/-^ mice were anesthetized by intraperitoneal injection of sodium pentobarbital(0.36 mg/10 g body weight) and xylazine(0.054 mg/10 g body weight). The abdomen skin was disinfected and subject to open abdomen surgery. Warm D-HBSS (no calcium, magnesium and phenol red) containing EDTA (0.5mM) and HEPES (25mM) was injected into liver through the vena cava and the portal vein was cut to let out blood. The portal vein was clamped with forceps when the liver was fully filled with D-HBSS buffer. The clamp and relaxing procedure was repeated until blood in the liver was completely washed out. Following the operation, warm digestion buffer(HBSS containing 1 mg/ml collagenase P and phenol red) was injected via the vena cava. The portal vein was clamped to make sure that digestion buffer completely filled the liver and was released to wash out digestion buffer. The clamp and relaxing procedure was repeated for 3-4 times. After digestion, the liver was removed and placed in a tube containing iced-cold HBSS. The liver was transferred into a 10-cm plate and the liver sack was ruptured with fine-tip forceps along the liver surface. After the rupture, the liver was gently pressed with cell scraper to release the cells and filtered through a 70 μm cell strainer into a 50ml tube. Iced-cold washing media (10 ml DMEM plus 10%FBS and 1% PS) was added and subject to centrifuge at 50 x g for 2 min. The supernatant was discarded and the pellets were resuspended with washing media. The procedure was repeated twice. Finally, the isolated hepatocytes were resuspended with plating media (DMEM plus 15%FBS and 1% PS) and plated in 6-well plates at the density of 4 x 10^5^ cells/per well. Plating media were changed after being cultured for 16 h.

When hepatocytes were cultured for 36-48 h, the media were switched to DMEM supplemented with1% FBS and 1% PS for 6 h, and further switched into DMEM supplemented with insulin (0.5μM), 1%FBS and 1% PS for 24h. During the period, L-ABH (125 μM) was added so that the hepatocytes were treated by L-ABH for 6 and 24h, respectively, before harvest. The cultures treated by saline were used for control.

## Hepatic glucose production

According to previous protocol [36], WT and *Srr*^-/-^ mice at the age of 8-12 weeks were used for primary hepatocyte cultures. The primary hepatocyte cultures were maintained for 72 h in DMEM with 15% FBS supplemented with 1% PS. The cultures were washed and maintained in Krebs-Ringer HEPES (KRH) buffer to fast cells for 2 h. The cells were further maintained in KRH buffer supplemented with sodium pyruvate(10 mM) for 6 h,either added with glucagon (100 nM),or with glucagon (100 nM) and L-ABH (125 μM). For treatments containing L-ABH, L-ABH (125 μM) was added 24 h before being washed with KRH buffer. The supernatant was collected for glucose analysis with the glucose assay kit (Solarbio),and normalized to the cellular protein contents.

## RT-qPCR

The liver was cut into pieces and homogenized in TRizol reagent (cat#15596-018, Invitrogen). RNA was dissolved in RNase-free ddH2O and the concentration was measured by use of a nanodrop 2000 microvolume spectrophotometer (ThermoFisher Scientific). Elimination of genomic DNA and synthesis of cDNA was performed by a first stand cDNA synthesis kit (cat# R111-02, Vazyme, NanJing, China). Fifty nanograms of cDNA were used as a template to amplify individual mRNA. The following primers were used in quantitative real-time polymerase chain reaction (RT-qPCR): m-*pepck*, sense, 5’-ACAGTCATCATCACCCAAGAGC-3’, antisense, 5’-CATAGGGCGAGTCTGTCAGTTC-3’; m-*g6pase*, sense, 5’-TCTTTCCCATCTGGTTCCATCT-3’, antisense, 5′- AATACGGGCGTTGT CCAAAC-3′; m-*rpl-19*, sense, 5′-CTCG TTGCCGGAAAAACA-3′, anti-sense, 5′- TCATCCAGGTCACCTTCTCA-3′.

The power SYBR™ Green PCR master mixture (cat#4367659, Invitrogen) was used for the qRT-PCR reaction following the manufacturer’s instruction. The amplification processes were initiated at 50 °C for 2 min and a secondary step at 95 °C for 10 min, followed by 40 cycles of PCR reactions (95 °C for 10 s and 60 °C for 1 min). Reaction specificity was examined by use of dissociative curves in which a single amplification peak for PCR reaction was present. The delta-delta Ct method was used to quantify the mRNA contents.

## Western blot

The liver tissues were washed, dissolved in Super RIPA buffer containing 1 x protease/1 x phosphatase inhibitor cocktail and protease cocktail (Sigma).

The tissues were homogenized, sonicated, and subject to centrifuge at 12,000 x g for 10 min. The supernatants were collected and resolved in 12% SDS- PAGE gels (Bio-Rad) and transferred to nitrocellulose membranes (ThermoFisher Scientific). Membranes were blocked in 5% skimmed milk and then incubated with primary antibody O/N at 4 °C. Following wash, the membranes were then incubated with secondary antibody at room temperature for 1 h, and developed with chemiluminescence system kit (Thermo Fisher Scientific). The blots were quantified using ImageJ software (NIH, Bethesda, MD, USA).

### *In vivo* toxicity test of L-ABH

Adult mice (C57BL/6) at the age of 8 weeks were fed water or L-ABH(20 mg/kg/d) until the age of 20 weeks. After fasting overnight, blood were collected from eye socket under anesthetization for examining the concentrations of aspartate aminotransferase, alanine aminotransferase, cystatin C, and insulin. The livers and kidneys were isolated and processed to H&E staining. The mice were immediately killed by detachment of cervical spine. The images were captured under phase-contrast microscope.

## Statistics

All data were indicated as average +/- SE. Sample sizes were chosen according to literatures and no statistical method was used to estimate sample size. All data were included in the study except decreased animals and thus no outlier was tested. The study was not blinded. Animals were assigned to different treatments with simple randomization. Data were examined with shapiro-Wilk for normality distribution. If data display normality distribution, the comparisons between two groups were performed with Student’s t-test and the differences among multiple groups were analyzed with one-way ANOVA with Tukey’s post hoc test. If data display non-normal distribution, the differences between two groups were performed with Mann-Whitney U test and those among multiple groups were analyzed with Kruskall-Wallis test (SPSS15.0.1; SPSS, Inc., Chicago, IL). Differences were considered significant if p<0.05.

Prism 6(GraphPad Software, Inc.), Image J (NIH), and Excel (Microsoft Corporation) were used for analyses.

## Data and Resource Availability

All the data supporting the results were reported in the main document and supplemented materials. Data sharing and resource sharing under appropriate request are available through the corresponding author.

## Results

### SRR was increased in the retinae of db/db mice

We have demonstrated the promoting role of SRR in DR [29, 30, 32, 33]. The role of SRR in DR was also recapitulated by an independent group [37]. In this study, we aim to investigate the therapentic potential by using an inhibitor of SRR in db/db mice, a T2DM model. First, we examined SRR expression in the retina and determined that SRR levels were significantly increased in db/db retinae, compared with those in WT mice of the same genetic background, BKS (Figures.1a and 1b). Correspondingly, D-Ser contents in the aqueous humor (0.407 +/- 0.0614 μg/mL) of db/db mice were significantly higher than those in WT mice at the values, 0.196 +/- 0.0138 μg/mL (p<0.05) (Figures. 1c-1f).

**Figure 1.**
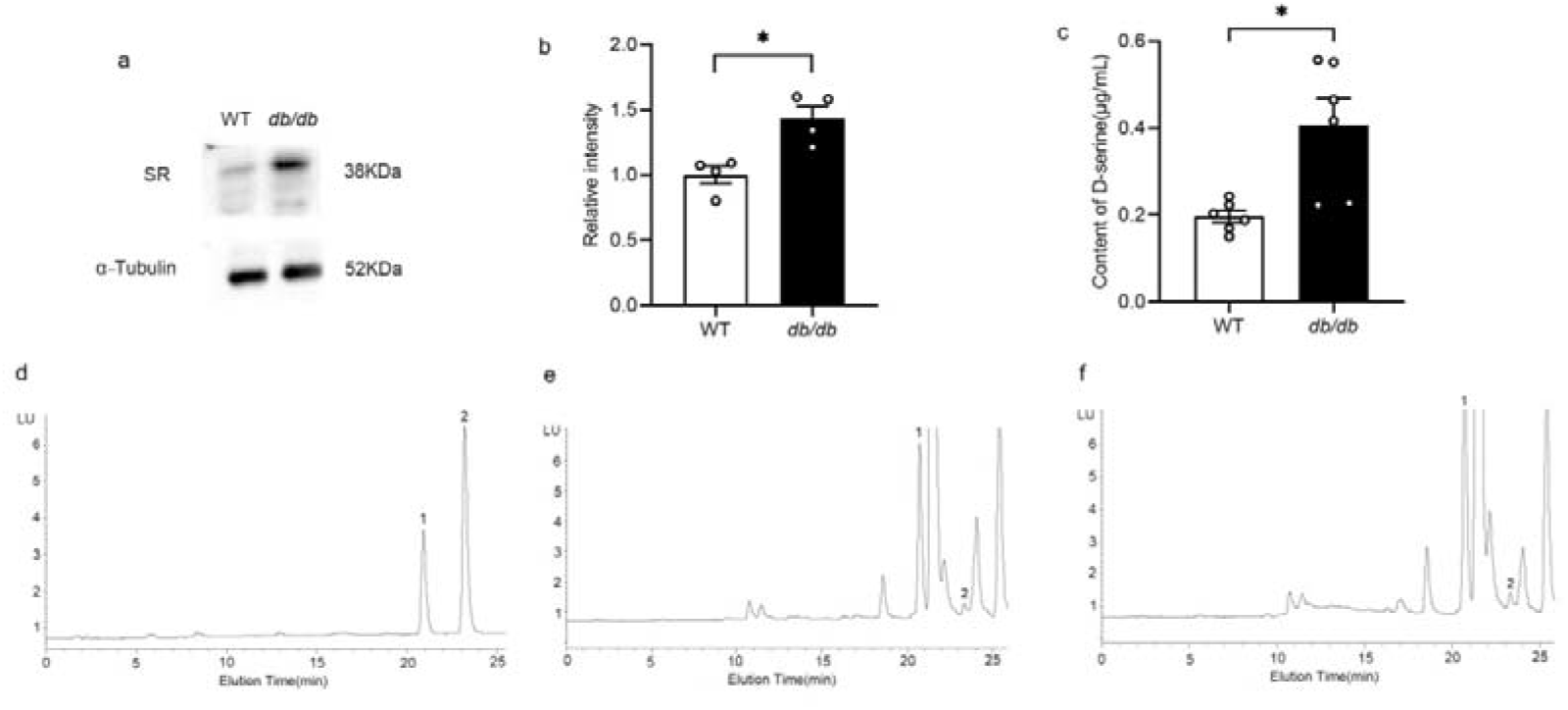
**SRR was increased in the retinae of db/db mice. (a)**The retinae of Db/db and WT mice with BKS background at the age of 7 months were homogenized and subjected to WB detection of SRR. (**b**) SRR levels in the retina were averaged from 4 retinae of db/d or WT mice. p<0.05. (**c**) D-Ser in aqueous humor from db/db and WT mice were determined with rp-HPLC. The contents were averaged from 6 mice of db/db or WT mice. p<0.05. (**d**) Amino acid standards were separated by rp-HPLC; 1: L-Ser, T_R_ = 20.511 min; 2: D-Ser, T_R_=23.284. Aqueous humor samples from WT (**e**) or db/db (**f**) mice. Student’s *t-test* was used to compare difference.

## Inhibition of SRR by oral gavage of L-ABH mitigates photoreceptor dysfunction in db/db mice

L-ABH is a selective and competitive inhibitor of SRR, forming an aldimine species with pyridoxal phosphate with an inhibition constant (Ki) at 97.5 +/- 23.7 μM [38]. Db/db male mice, at the age of 12 weeks, were fed by oral gavage of L-ABH (20 mg/kg/d) or equal amount of water for 18 weeks. ERG recordings were conducted to evaluate photoreceptor and phototransduction function after 18 weeks of feeding. Under scotopic condition, the b-wave deficits in db/db mice were rescued by L-ABH. In particular, under 0.01cd-s/m^2^stimulus intensity, the b-wave amplitudes in db/db mice were 19.4 +/- 5.49 μV, significantly less than 88.5 +/- 19.86 μV in WT mice (p<0.01); L-ABH administration increased the amplitude in db/db mice to 82.6 +/- 12.09 μV (p<0.01) (Figure.2a). Under 3 cd-s/cm2 stimulus intensity, the b-wave amplitudes in db/db mice were 42.3+/- 9.67 μV, significantly less than 184.8 +/- 29.27 μV (p<0.01); L-ABH administration increased the amplitudes in db/db mice to 128.2 +/- 26.89 μV (p<0.05) (Figure.2b). Under 10 cd-s/cm2 stimulus intensity, the b-wave amplitudes in db/db mice were 45.5 +/- 11.93 μV, significantly less than 206.6 +/- 44.26 μV (p<0.01); L-ABH administration increased the amplitudes in db/db mice to 143.8 +/- 27.54 μV (p<0.05) (Figure. 2c). The oscillatory potentials in db/db mice were also rescued by L-ABH. For example, the amplitudes of OS1, OS2, OS3 in db/db mice were 32.1 +/- 9.36 μV, 21.2 +/- 9.13 μV, 23 +/- 7.66 μV, significantly less than 122.5 +/- 30.85 μV (p<0.05), 134.6 +/- 40.13 μV (p<0.05), 152.8 +/- 45.52 μV (p<0.01), respectively. L-ABH increased the amplitudes of OS2 and OS3 in db/db mice to 101.6 +/- 28.50 μV (p<0.05), 118.1+/- 19.06 μV (p<0.01), respectively whereas L-ABH did not increase OS1 amplitude to significant levels, at 103.4 +/- 20.10 μV (n.s vs. db/db mice + water) (Figure.2d).

**Figure 2.**
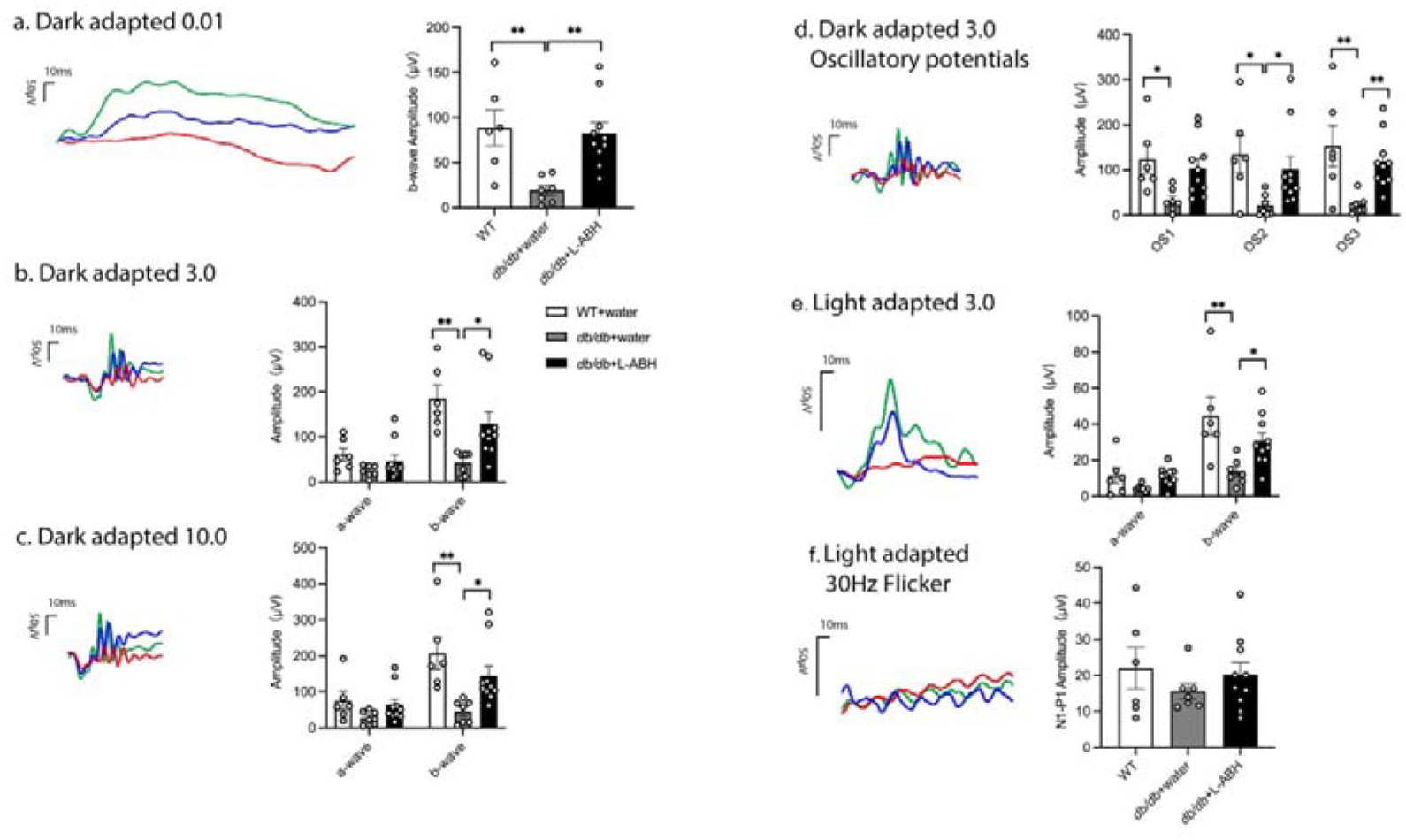
The effects of L-ABH on photoreceptor function. Db/db male mice were fed for 18 weeks with water (db/db+water) or L-ABH (20 mg/kg/d) (db/db+L-ABH). BKS WT mice on regular chow diet were included (WT). At the end of feeding, the mice were subjected to ERG recording. Under scotopic condition, a and b-wave were recorded under 0.01 (**a**), 3 (**b**), 10 (**c**) cd-s/m^2^ stimuli. (**d**) Under 3 cd-s/m^2^, oscillatory potentials were recorded. (**e**) Under photopic condition, the amplitudes for a and b-wave were recorded under 3 cd-s/m^2^ stimulus intensity. (**f**) Under photopic condition, the responses of flicker were recorded under 30 Hz flicker light. The amplitudes were averaged from 6-10 mice. *p<0.05, **p<0.01, Kruskall-Wallis test was used to compare the differences.

Under photopic condition, 3 cd-s/cm2 stimulus intensity, the b-wave amplitudes in db/db mice were 13.5 +/- 2.68 μV, significantly less than 44.4 +/-10.39 μV (p<0.01); L-ABH administration increased the amplitudes in db/db mice to 30.6 +/- 4.46 μV (p<0.05) (Figure.2e). Under photopic condition, 30 Hz flicker was used to stimulate the retina under which there were no differences among the three groups with respect to the amplitudes of N1 and P1 (n.s) (Figure.2f).

## Inhibition of SRR by oral gavage of L-ABH mitigates neurovascular pathologies in db/db mice

To evaluate retinal thickness, circular and linear scans with SD-OCT examination were conducted. Under circular mode,the retina was divided to four quadrants and the measurement focused on the thickness of RGCL (Figure.3a). In the nasal, ventral, and temporal retina of db/db mice, the thickness were 18.3 +/- 0.91 μm, 11.6 +/- 2.97 μm, 14.2 +/- 1.25 μm, significantly less than 22.8 +/- 1.26 μm (p<0.05), 24.8 +/- 0.96 μm (p<0.01), 19.7 +/- 1.02 μm (p<0.05) in the individual quadrant of WT retina. L-ABH increased the nasal retinal of db/db mice to significances, 22.3 +/- 0.63 (p<0.05 vs. db/db + water) whereas the increases were not different for the ventral and temporal retina (n.s, respectively) (Figure.3b). In dorsal retina, there were no differences regarding the retinal thickness among the three groups (Figure.3b). We further analyzed retinal thickness with linear scan in which the thickness of each layer of the retina was determined (Figure.3c). At the nasal side, the retina at 200-1400 μm from optic nerve head (ONH) in db/db mice were much thinner than WT mice and L-ABH significantly increased the thickness in db/db mice to the similar levels in WT mice (p<0.05). By contrast, the protection of L-ABH on the temporal side of ONH did not achieve significance (Figure.3d). RNFL-RGCL-IPL in db/db mice at 200-500 μm from the temporal side of ONH was much thinner than WT mice and L-ABH significantly increased thickness at these regions (p<0.05, 0.001) (Figure.3e). By contrast, L-ABH did not significantly increase the thickness of RNFL-RGCL-IPL from the nasal side of ONH (Figure.3e). Regarding the thickness of INL in db/db mice, L-ABH significantly increased retinal thickness at 700-1300 μm of the temporal side and those at 400-1400 μm of the nasal side of ONH (p<0.05,0.01)(Figure.3f). Regarding the thickness of OPL in db/db mice, L-ABH significantly increased retinal thickness at 400-1100 μm of the nasal side and those at 1000-1500 μm of the temporal side of ONH (p<0.05,0.01) (Figure.3g). In outer retina, there were no difference among three groups (Supplemental Fig.1). In conclusion, L-ABH protected inner retina.

**Figure 3.**
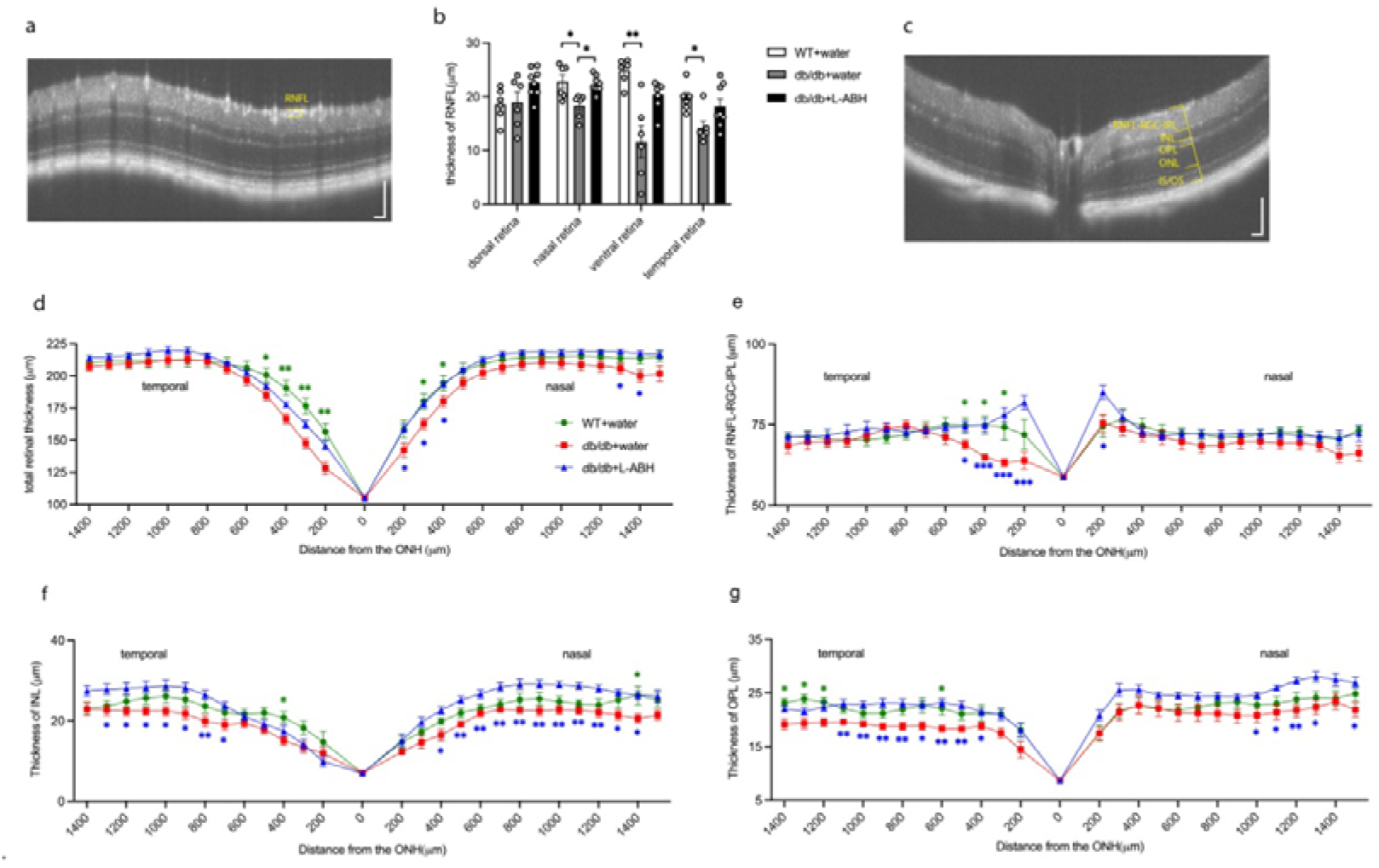
The effects of L-ABH on the structure of neuroretina. Db/db male mice were fed for 18 weeks with water (db/db+water) or L-ABH (20 mg/kg/d) (db/db+L-ABH). BKS WT mice on regular chow diet were included (WT). At the end of feeding, the mice were subjected to SD-OCT examination. (**a**) Under circular mode of scan, the optic disk was wrapped by the scan box and the retina surrounding optic disk was automatically scanned. (**b**)The RNFL thickness of each retina was quantified for the dorsal, nasal, ventral, and temporal quadrant. (**c**) Under linear mode of scan, the optic disk was positioned in the center of the scan box and the images were automatically acquired from the nasal to the temporal retina. The total thickness from RNFL to IS/OS (**d**), of RNFL-RGC-IPL (**e**), INL (**f**), OPL (**g**) were recorded under 200 μm interval with reference to optic nerve head (ONH) as the central point. The quantifications were averaged from 6-10 mice. *p<0.05, **p<0.01. * p<0.05, **p<0.01 indicated differences between db/db +water and WT +water. *p<0.05, **p<0.01, ***p<0.001 indicated differences between db/db + L-ABH and db/db + water. Kruskall-Wallis test was used to compare the differences.

We further evaluated RGC density in the retina among the three groups. With retinal flatmounts, the RGCs were labelled with a specific nuclear marker, Brn3a. In the central retina, there were no difference of RGC densities among the groups of WT, db/db mice fed with water, or with L-ABH (Figure.4a). In the intermediate retina, RGC densities in db/db mice were 1500 +/- 66.70 /mm^2^, significantly less than 1693.8+/- 61.39/mm^2^ in WT mice (p<0.05); with L-ABH administration in db/db mice, the densities were increased to 1833.9 +/-39.61/mm^2^ (p<0.01) (Figure.4b). In the peripheral retina, RGC densities in db/db mice were 1062 +/- 53.20 /mm^2^, similar as 1195.9 +/- 65.60/mm^2^ in WT mice; with L-ABH administration in db/db mice, the densities in db/db retinae were increased to 1250.4 +/- 46.82/mm^2^ (p<0.05) (Figure.4c). Thus, L-ABH protected against retinal neurodegeneration in db/db mice.

**Figure 4.**
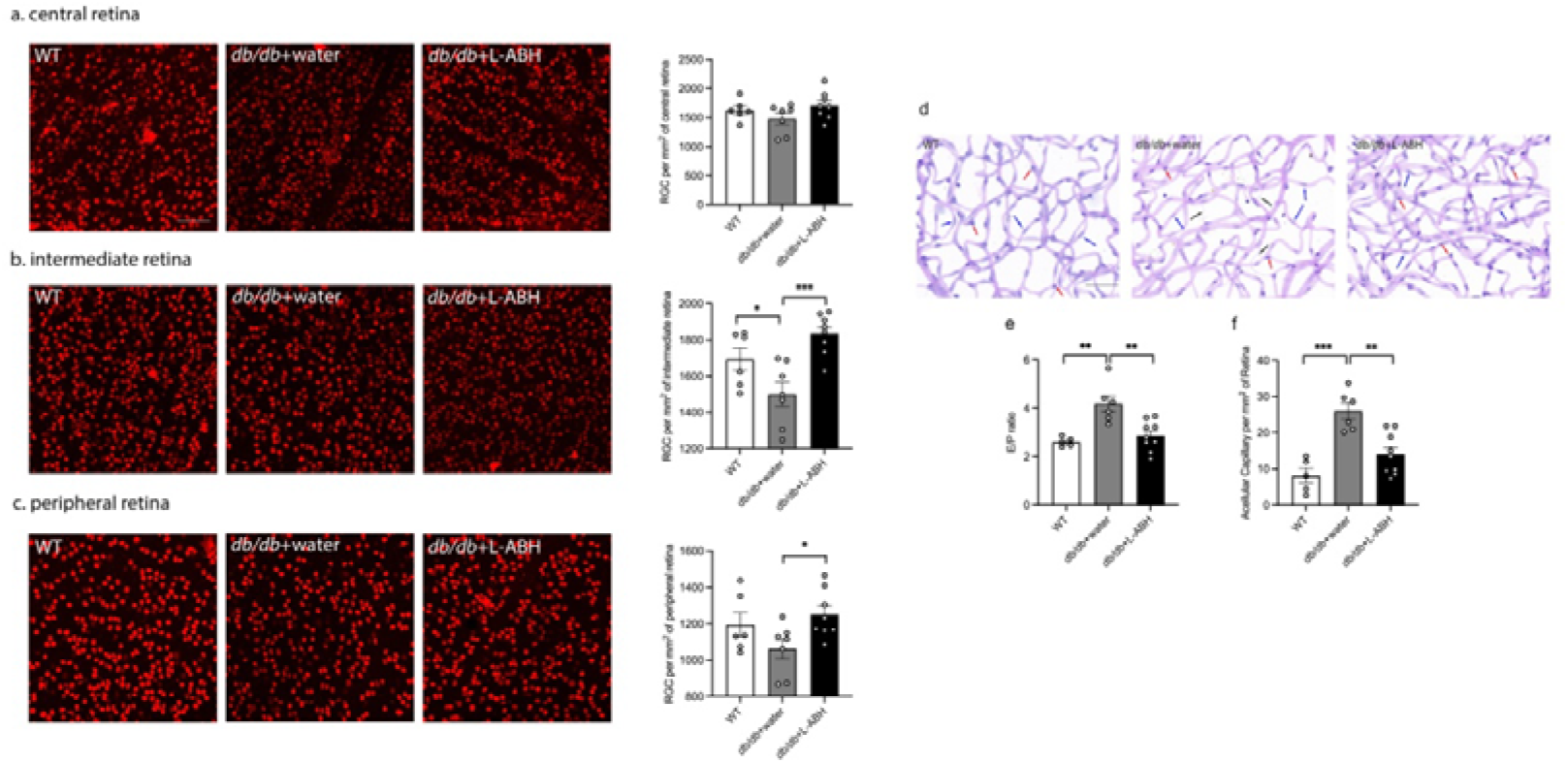
The effects of L-ABH on RGC. Db/db male mice were fed for 18 weeks with water (db/db+water), L-ABH (20 mg/kg/d)(db/db+L-ABH). BKS WT mice on regular chow diet were included (WT). At the end of feeding, the retina were made into flatmounts and subjected to staining with Brn3a. The RGC numbers were quantified for the central (**a**), intermediate (**b**), and peripheral retina (**c**), from 6-8 mice. (**d**) Some of the retinae were subjected to trypsin digestion, followed by PAS staining and haematoxylin counterstaining. Red arrow indicated pericytes; blue, endothelial cells; dark, acellular capillary. The ratios between endothelial cells and pericytes (E/P) (**e**) and the numbers of acellular capillary (**f**) were quantified from 6-8 mice. *p<0.05, **p<0.01, ***p<0.001, Kruskall-Wallis test was used to compare the differences.

We further evaluated whether inhibition of SRR by oral gavage of L-ABH impacted retinal microvessels in db/db mice. With periodic acid-Schiff stain and heamatoxylin counterstaining, retinal vasculature was indicated with clear images of retinal endothelial cells and pericytes (Figure.4d). The ratios of endothelial cells and pericytes in diabetic retina were 4.2 +/- 0.33, significantly higher than 2.6 +/- 0.09 in WT mice (p<0.01); L-ABH normalized the ratios to 2.9 +/- 0.20 (p<0.01) (Figure.4e). We also quantifed the numbers of avascular capillary. The numbers of acellular capillary in db/db mice were 26.0 +/- 2.22/ mm^2^, significantly higher than 8.1 +/- 2.07/mm^2^ in WT mice (p<0.001); L-ABH significantly reduced the numbers to 14.0 +/- 1.90 /mm^2^ (p<0.01) (Figure.4f).

## Intravitreal injection of L-ABH protects against glutamate-induced neurotoxicity

Glutamate-induced neurotoxicity contributes to RGC degeneration in diabetic retina [24–26]. We thus investigated whether the neuroprotection provided by L-ABH was in part through antagonizing glutamate-mediated neurotoxicity. Intravitreal injection of glutamate (1 μL,100 mM) significantly induced apoptosis, mostly in RGCL and INL (Figure.5a). To model the concentration of circulating level of L-ABH by oral gavage, L-ABH (1 mg/mL) was intravitreally injected along with glutamate. L-ABH significantly reduced RGC apoptosis (Figure.5a). To quantify the numbers of Brn3a + cells for TUNEL staining, the numbers in the group of glutamate injection were 30 +/- 2/mm^2^, significantly higher than the injection of L-ABH plus glutamate at the values, 14 +/- 2/mm^2^ (p<0.001). There were no apoptosis in the retina of WT mice subjected to saline injection (Figure.5b).

**Figure 5.**
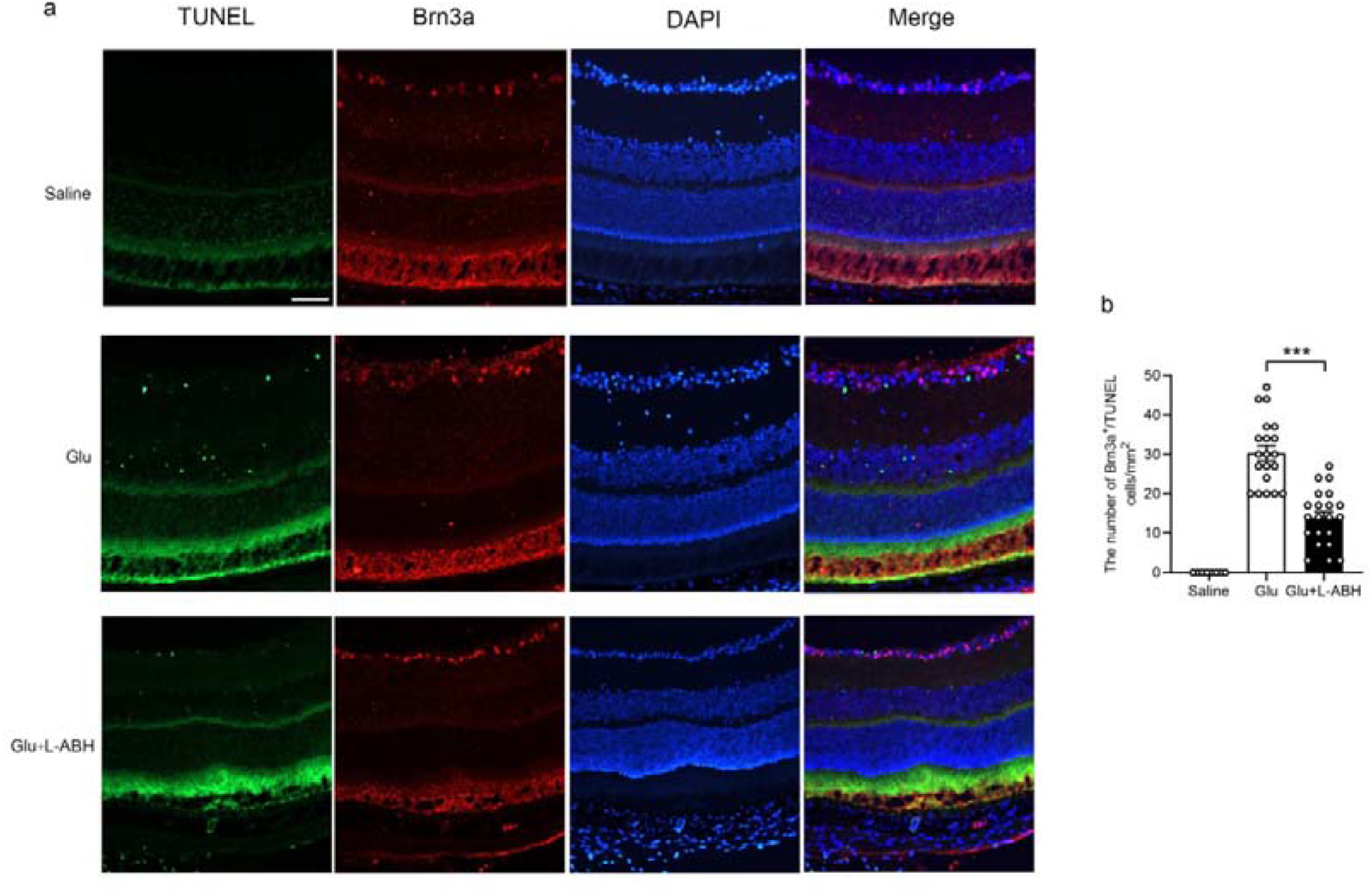
The effect of intravitreal injection of L-ABH on glutamate-induced RGC apoptosis. (a) C57BL/6J mice at the age of 2 months were subjected to intravitreal injection of 1 μL of saline, glutamate (100 mM), or glutamate plus L-ABH (1 mg/mL). After 24 h, the retina were made into cryosections and subjected to TUNEL and Brn3a staining. At the end of staining, DAPI was used to stain the cell nuclei. (b) The RGC cells positive for TUNEL staining were quantified from 2 mice for saline, 4 mice for either glutamate or glutamate plus L-ABH injection, 3-4 sections for each eyeball. ***p<0.001, Kruskall-Wallis test was used to compare the differences.

## Inhibition of SRR by L-ABH improves glucose intolerance and mitigates insulin resistance in db/db mice

Since glycemic control reduces the incidence and the progression of DR. We wonder whether the protective effects of L-ABH on the neurovascular abnormalities in db/db mice are partly mediated via glycemic normalization. Previous studies suggest that single-nucleotide polymorphism of *srr* is associated with T2DM [39, 40]. Further study demonstrated that mice with whole body deletion of *srr* show improved glucose tolerance through enhanced insulin secretory capacity [41]. Thus, We further investigated whether inhibiton of SRR impacted glucose homeostasis. Similarly as the above procedures, db/db male mice, at the age of 12 weeks, were fed L-ABH (20 mg/kg/d) or equal amount of water for 18 weeks. Simultaneously, metformin feeding (250 mg/kg/d) was used as a positive control. In parallel, WT mice with BKS background were assigned to compare the efficacy of treatments. L-ABH did not alter body weight gain compared with vehicle feeding in db/db mice whereas metformin did (Figure.6a). At two weeks after L-ABH feeding, random blood glucose significantly decreased relative to water feeding (**p<0.01), in db/db mice (Figure.6b). After 18 weeks of L-ABH feeding, the random blood glucose levels in db/db mice reached the levels at 135.9+/-11.86 mg/dL, similar to the values in WT mice under normal chow diet (Figure.6b). At the end of feeding, db/db mice fed with L-ABH displayed improved glucose tolerance (AUC under L-ABH, 31809.7+/-1553.87 mg.dL^-1^.min) compared with water feeding (AUC under water feeding, 59757+/-1676.62 mg.dL^-1^.min) (***p<0.001). Similarly, feeding metformin improved glucose tolerance (AUC, 40579.5+/- 2869.97 mg.dL^-1^.min)(*p=0.022 *vs*.water)(Figures.6c and 6d).

**Figure 6.**
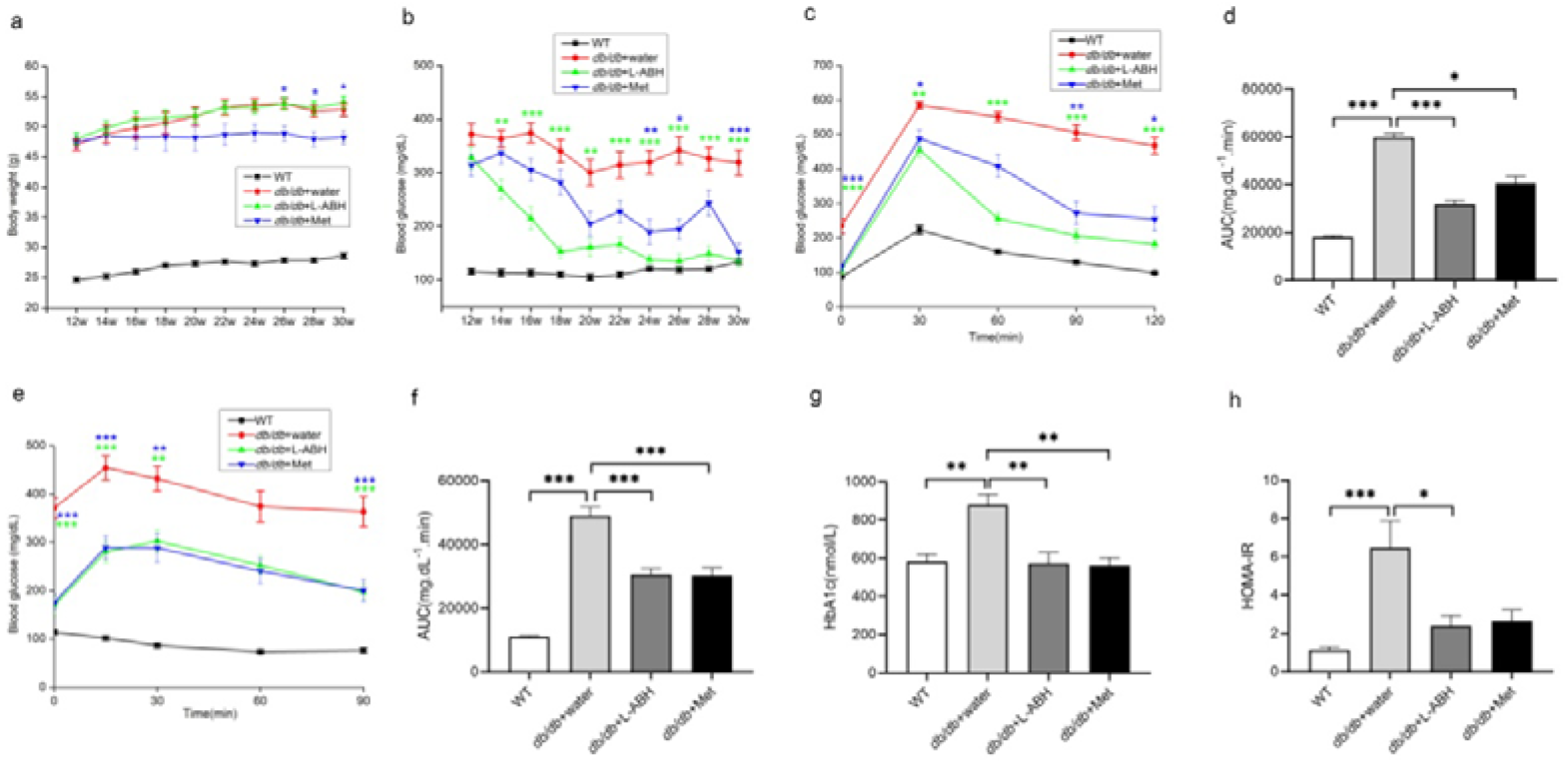
The effects of L-ABH feeding on glucose homeostasis and insulin sensitivity in db/db mice. Db/db male mice were fed for 18 weeks with water(db/db+water), L-ABH(20 mg/kg/d)(db/db+L-ABH), metformin(250 mg/kg/d)(db/db+Met) and BKS WT mice on regular chow diet were also included (WT). (**a**)The body weights for WT(n=14), db/db+water(n=25), db/db+L-ABH (n=22), db/db+metformin(n=19). db/db+Met *vs*.db/db+water, *p<0.05. (**b**)The random blood glucose for WT (n=14), db/db +water(n=25), db/db+L-ABH(n=22), db/db+metformin(n=19). **p<0.01, ***p<0.001 indicated significant differences between db/db+L-ABH and db/db+ water at indicated time points. *p<0.05, **p<0.01, and ***p<0.001 indicated differences between db/db+Met and db/db+water. (**c**) OGTT for WT(n=18), db/db+water(n=21), db/db+L-ABH(n=22), db/db+ metformin(n=18). ***p<0.001 indicated differences between db/db+L-ABH and db/db+water. *p<0.05, **p<0.01, and ***p<0.001 indicated differences between db/db+Met and db/db+water. (**d**)Areas under curve for OGTT in **c**. (**e**)ITT for WT(n=16), db/db+water(n=21), db/db+L-ABH(n=22), db/db+metformin(n=18). **p<0.01, ***p<0.001 indicated differences between db/db+ L-ABH and db/db+ water. **p<0.01,***p<0.001 indicated differences between db/db+Met and db/db+water. (**f**) Areas under curve for ITT in **e**. (**g**) HbA1c for WT(n=10), db/db+water(n=8), db/db+L-ABH (n=10),and db/db+metformin (n=6). (**h**)HOMA-IR values for WT(n=14),db/db+ water(n=15), db/db+L-ABH(n=10),and db/db+ metformin (n=9). One-way ANOVA with repeated measures for inter-treatment evaluations at each time point(**a,b,c,e**). One-way ANOVA for inter-treatment evaluations(**f,g**) and Kruskall-Wallis for evaluating differences(**d and h**). All the results were averaged from triplicate experiments

L-ABH feeding increased insulin sensitivity (AUC, 30560.3+/- 1930.72 mg.dL^-1^.min) compared with water feeding (48857.1+/- 2906.24 mg.dL^-1^.min)(***p<0.001). Similarly, feeding metformin increased insulin sensitivity (30187.5+/- 2536.96 mg.dL^-1^.min)(***p<0.001 *vs*.water)(Figures.6e and 6f). By monitoring HA1C, the levels in WT BKS mice were 582.1+/-37.37 nmol/L (Figure.6g). The levels of HA1C treated by L-ABH in db/db mice reached 572.4+/-56.29 nmol/L which were significantly lower than the values under water feeding (877.9+/-56.19 nmol/L) (**p=0.009). Similarly, metformin reduced HA1C in db/db mice to 561.2+/-40.31 in db/db mice (**p=0.004 *vs*.water)(Figure.6g). Regarding HOMA-IR values, L-ABH significantly reduced HOMA-IR at the values, 2.4+/-0.55, compared with the values at 6.5+/-1.40 under water feeding (*p=0.046) whereas metformin reduced HOMA-IR to 2.6 +/-0.61 which were not significantly different with the values from water feeding (p=0.167)(Figure.6h).

Previous study indicated that whole-body deletion of SR increases *in vivo* glucose stimulated insulin secretion(GSIS) [41]. Thus, it suggests that L-ABH may modify GSIS which contributes to imporved glucose tolerance and insulin sensitivity. We examined whether L-ABH modified GSIS in isolated pancreatic islet cultures. Insulin secretion was indicated by the ratios between insulin levels in the supernatants and the protein content in the culture well. The ratios under low glucose condition(Con+LG) were 9.62 x 10^-4^ +/- 2.33 x 10^-4^, compared with the values at 1.37 x 10^-3^ +/- 4.91 x 10^-4^, under simultaneous additon of L-ABH(125 μM) with low glucose (L-ABH+LG) (p=0.471) (Supplemental Fig.2a). The ratios under high glucose (Con+HG) were 1.15 x 10^-3^ +/- 2.34 x 10^-4^, compared with the values at 1.05 x 10^-3^ +/- 3.171 x 10^-4^, under simultaneous addition of L-ABH (125 μM) with high glucose (L-ABH+HG) (p=0.795) (Supplemental Fig.2a). Under both low and high glucose condition, L-ABH did not change the amount of supernatant insulin compared with vehicle treatment in pancreatic islet cultures. That the insulin secretion was unaltered by L-ABH in pancreatic islets was consistent with the observation:unaltered blood insulin levels in db/db mice, under L-ABH feeding, relative to water feeding(Supplemental Fig.2b). The blood insulin levels were: 4.4+/-0.58 mIU/L for WT BKS; 7.0+/-1.18 mIU/L for db/db mice fed with water; 3.8+/-0.77 mIU/L for db/db mice fed with L-ABH; 5.2+/-1.00 mIU/L for db/db mice fed with metformin(Supplemental Fig2b).

By using CLAMS cages, metabolism of mice were monitored including consumption of water and food, energy expenditure and locomotor activity. Under L-ABH feeding, water consumption in db/db mice was substantially reduced compared with db/db mice under water feeding (Supplemental Fig.3a). By contrast, food consumption in db/db mice under L-ABH feeding remained similar as db/db mice under water feeding (Supplemental Fig.3b). L-ABH modestly increased locomotor activity in db/db mice compared with water feeding under light cycle and the increased amplitude was similar to metformin (Supplemental Fig.3c). L-ABH or metformin feeding did not alter respiratory quotient relative to water feeding, although the oxygen consumption and production of carbon dioxide in db/db mice were significantly lower than WT BKS mice. The similar respiratory quotient suggests that energy expenditure remained at the same levels between L-ABH and water feeding of db/db mice(Supplemental Figs.3d-3f). To test the efficacy of L-ABH to inhibit SRR, serum l-/D-serine levels were measured with reverse-phase HPLC which indicated that l-serine and l-/D-serine ratios were modestly increased by L-ABH feeding relative to water feeding, respectively(Supplemental Fig.4).

Thus, SRR was effectively inhibited by ingestion of L-ABH.

## L-ABH or genetic deletion of *Srr* improves glucose tolerance and insulin sensitivity in DIO mice

To confirm whether the anti-diabetic effects by L-ABH were via specific inhibition on SRR, we initially planned to crossbreed *Srr*^-/-^ mice with db/db mice. However, db/db mice were created on BKS background and *Srr*^-/-^ mice generated in our recent work were established on C57BL/6 species [35].

Diabetic phenotypes can be recapitulated on BKS but not on C57BL/6 background. Thus,we gave up the original plan and compared the effects of L-ABH and *Srr* deletion on DIO mice.

To investigate SRR inhibition on glucose homeostasis, C57BL/6 WT male mice were fed HFD for 16 weeks since the age at 3 weeks. On the third day after HFD, the mice were subject to oral gavage of L-ABH (20 mg/kg), or of water. Different with the effect on db/db mice, L-ABH reduced weight gain under HFD (35.4 +/- 1.57g), relative to water feeding(41+/-0.83g), at the end of feeding (**p=0.008), aged 19 weeks (Figure.7a). At the ages of 18 and 19 weeks, L-ABH reduced random blood glucose levels relative to water feeding (*p<0.05, respectively) (Figure.7b). At 12 weeks of feeding, the mice were subject to insulin sensitivity assay. L-ABH feeding improved ITT(AUC, 10206.3 +/- 274.01 mg.dL^-1^.min), compared with water feeding (11187.9+/- 208.85 mg.dL^-1^.min) (*p=0.015), in WT DIO mice (Figures.7c and 7d). The mice were continued to be fed for 4 more weeks and OGTT was tested. Under HFD, L-ABH feeding improved glucose tolerance (AUC, 28749.43 +/- 1785.45 mg.dL^-1^.min), compared with water feeding (37937.0+/- 1690.79 mg.dL^-1^.min) (**p=0.003), in WT DIO mice (Figures.7e and 7f). In paralle, we tested *Srr* deletion on glucose tolerance and insulin sensitivity in *Srr*^-/-^ DIO mice. *Srr*^-/-^ and littermate WT mice with C57BL/6 background, at the age of 3 weeks, were fed with HFD for 16 weeks. After 12 weeks of feeding in WT mice, HFD impaired insulin sensitivity (AUC, 9882.1+/-512.32 mg.dL^-1^.min) compared with under normal chow diet (7521.5+/- 472.56 mg.dL^-1^.min) (*p=0.011). Under HFD, *Srr*^-/-^ mice indicated similar insulin sensitivity (7310.5+/-699.15 mg.dL^-1^.min), as under normal chow diet (7224.7+/-792.38 mg.dL^-1^.min)(p=0.924) (Figures.7g and 7h). After 12 weeks of feeding, the fasting blood glucose levels in *Srr*^-/-^mice under HFD were significantly lower than those in WT mice under HFD(**p<0.01) (Figure.7i). However, at the end of 16 weeks of feeding, the fasting blood glucose levels in *Srr*^-/-^ mice under HFD were similar as those in WT under HFD (Figure.7j). Regarding glucose tolerance, the values in *Srr*^-/-^ DIO mice (AUC, 27961.9+/- 1509.04 mg.dL^-1^.min) were significantly improved, compared with the values in WT DIO mice (36544.5+/-2030.41 mg.dL^-1^.min) (**p=0.002)(Figures.7k and 7l). Regarding OGTT and ITT assays, the amplitudes of alterations between L-ABH and water feeding in DIO mice were similar as the differences between WT mice and *Srr*^-/-^ mice under HFD. These observations suggest that the effects of L-ABH on glucose homeostasis were predominantly through inhibition of SRR.

**Figure 7.**
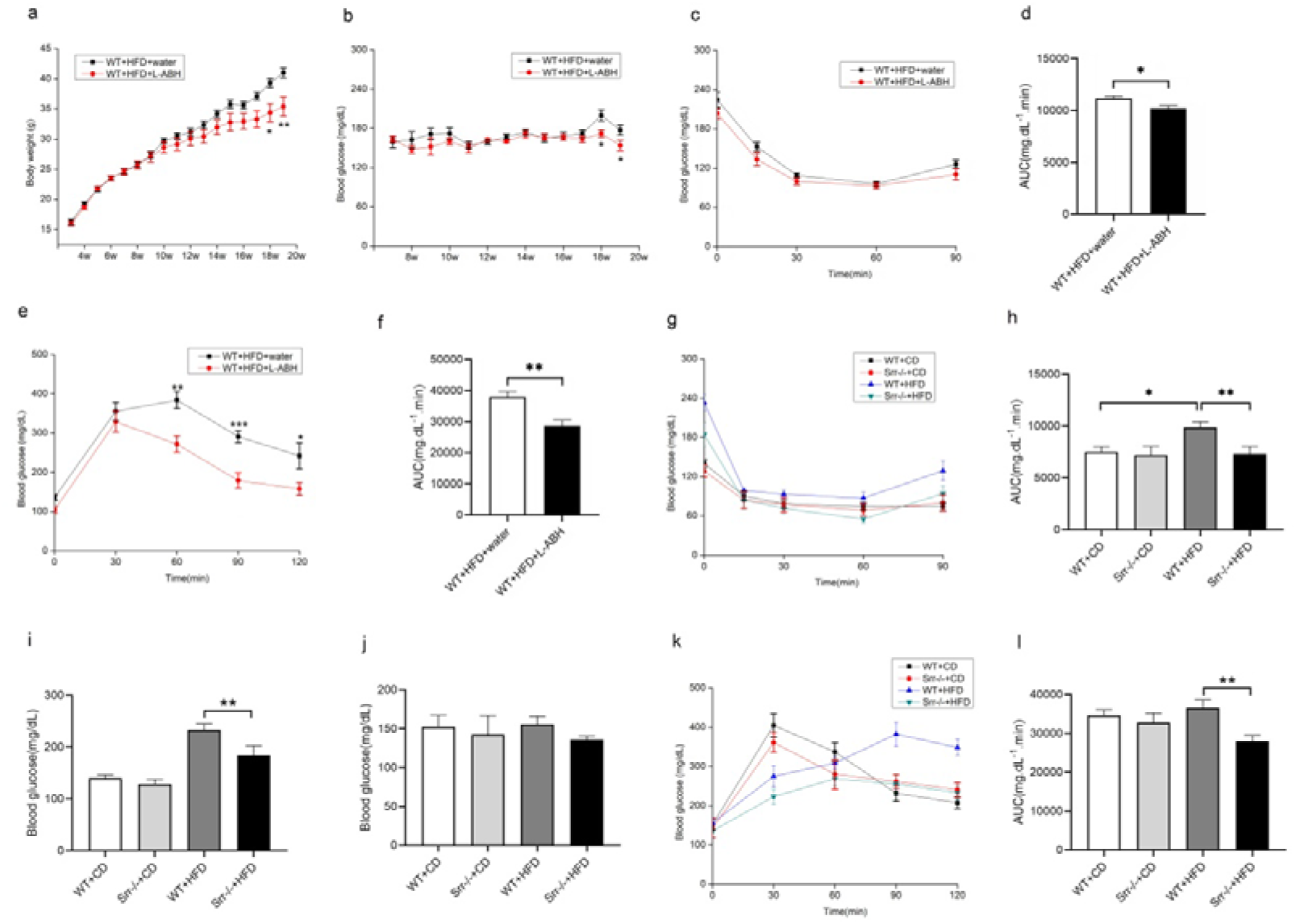
The effects of L-ABH feeding or *Srr* deletion on glucose homeostasis and insulin sensitivity in DIO mice. a-f:the effects of L-ABH feeding on DIO mice. WT male C57BL/6 mice were fed HFD for 16 weeks. (**a**)The body weight for L-ABH(WT+ HFD+L-ABH)(n=7) or water feeding(WT+HFD+water)(n=7). *p<0.05 and **p<0.01 indicated differences at the age of 18 weeks and 19 weeks,respectively. (**b**)The random blood glucose levels for L-ABH (n=7) or water feeding(n=7). *p<0.05 indicated significant differences at the age of 18 and 19 weeks, respectively. (**c**) At 12 weeks of feeding, ITT was conducted as aforementioned for DIO mice under L-ABH or water feeding. (**d**) Areas under curve for ITT in **c**. *p<0.05. (**e**) OGTT was performed as aforementioned at 16 weeks of feeding. (**f**) Areas under curve for OGTT in **e**. **p<0.01. g-l: the effects of *Srr* deletion on DIO mice. WT or *Srr*^-/-^ male C57BL/6 mice were fed HFD or normal chow(CD) for 16 weeks. (**g**)At the age of 15 weeks, ITT assay for WT(n=8),*Srr*^-/-^ +CD(n=7),WT +HFD (n=8),and *Srr*^-/-^ +HFD(n=8). (**h**) Areas under curve for ITT in **g**. **p<0.01. (**i**)The fasting blood glucose levels at the age of 15 weeks. *p<0.05. (**j**) The fasting blood glucose levels at the age of 19 weeks. (**k**) At the end of feeding, the mice were subject to OGTT assay. (**l**)Areas under curve for OGTT in **k**. **p<0.01. One-way ANOVA with repeated measure for the inter-treatment evaluations.

## L-ABH inhibits liver gluconeogenesis in db/db mice and deletion of *Srr* **reduces glucagon-induced glucose production in hepatocyte cultures**

To investigate the mechanism by which L-ABH normalized blood glucose and improved insulin sensitivity, we first examined SRR expression in the tissues involving in glucose metabolism including the liver, the pancreas, and the muscles. Regarding the amount of expression, the levels of SRR expression were ranked in the order: brain> liver> pancreas∼= gastrocnemius in C57BL/6 WT mice (highest expression in the brain)(Figure.8a). The glycogen levels were not different in the liver or in the gastrocnemius muscle of db/db mice between L-ABH and water feeding (Supplemental Figures.5a and 5b). We further examined the rate-limiting enzymes of liver gluconeogenesis including glucose-6- phosphatase (G6Pase) and phosphoenolpyruvate carboxykinase (PEPCK). In db/db mice feeding with L-ABH, the mRNA levels of *g6pase* and *pepck* in the liver were significantly reduced compared with those of db/db mice fed with water (Figures.8b and 8c). Correspondingly, the protein levels of G6Pase and PEPCK in the liver under L-ABH feeding were significantly reduced compared with water feeding in db/db mice (Figures.8d-8f). To examine whether the effect of L-ABH feeding on G6Pase and PEPCK was mediated via SRR, we tested *in vitro* with primary hepatocyte cultures derived from WT and *Srr*^-/-^ mice. Consistent with *in vivo* results, L-ABH reduced the levels of *pepck* and *g6pase* mRNAs compared with sham treatment in primary hepatocyte cultures from WT mice, under the context of high glucose and high insulin. In cultures derived from *Srr*^-/-^ mice, L-ABH did not alter the mRNA levels of *pepck* and *g6pase* compared with sham treatment (Figures.8g and 8h). The differential regulation by L-ABH in WT and *Srr*^-/-^ hepatocytes suggests that inhibition of gluconeogenesis by L-ABH is dependent on the presence of *srr*. Consistently, L-ABH modestly inhibited glucose production in primary hepatocyte cultures by glucagon, compared with sham treatment whereas the glucose production induced by glucagon was significantly reduced in *Srr*^-/-^ compared with WT hepatocyte cultures (Figure.8i).

**Figure 8.**
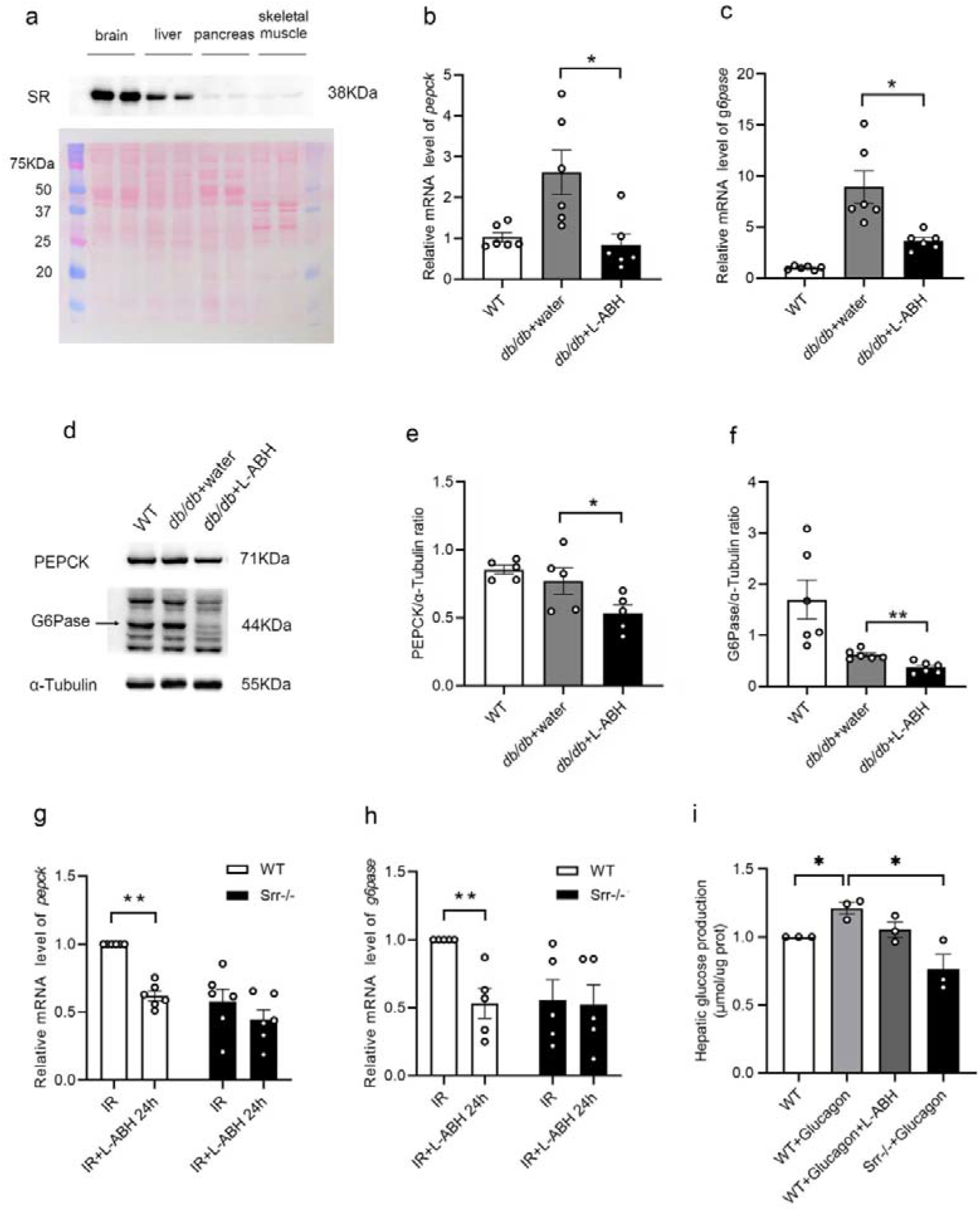
The inhibitory effects of L-ABH on hepatic *pepck and g6pase* was dependent on *Srr*. (**a**) **Top:**The cerebral cortex,the liver,the pancreas,the gastrocnemius muscle were isolated from WT C57BL/6 male mice(n=2) and harvested for western blot against SR. Equal amount of protein(40 μg) for each well was loaded. **Low**: Ponceau red staining. (**b and c**)The livers were isolated from db/db mice feeding with L-ABH(n=6), water(n=6), or WT(n=6) and subject to RT-qPCR examination of *pepck and g6pase* mRNA. (**d-f**)The livers were isolated from db/db mice feeding with L-ABH(n=5∼6), water(n=5∼6), or WT(n=5∼6) and harvested for WB examination of PEPCK and G6Pase. (**g and h**)The primary hepatocyte cultures from WT and *Srr*^-/-^ mice were maintained in DMEM plus 10% FBS until 50-60% confluency and switched to DMEM plus 1% FBS for 6 h. High glucose(25 mM) and insulin(0.5 μM)(IR) or L-ABH(125 μM) were simultaneously added to the cultures for 24 h. The cells were harvested for RT-qPCR. Mann-Whitney U was used to compare the difference between IR and IR plus L-ABH conditions in WT or *Srr*^-/-^ hepatocyte cultures. (**i**)The hepatic glucose production was compared among individual treatment. One-way ANOVA with repeated measures for inter-treatment evaluations (**c,e,f,i**) and Kruskall-Wallis for differences(**b**).*p<0.05,**p<0.01 indicated differences between indicated groups.

## L-ABH does not show toxicity towards the kidney and the liver

Notably, L-ABH and its D-isomer indicate anti-tumor activity with comparable efficacy and l-ABH,with LD50 in leukemia mice at 3,300 mg/kg by i.p injection, indicates higher toxicity than D-ABH in leukemia mice [42]. Thus, we tested its potential toxicity in WT C57BL/6 mice. After L-ABH feeding for 12 weeks at the dose of 20 mg/kg/d, the liver and kidney were extracted to examine morphology and blood samples were collected to examine the functions of the organs. L-ABH did not induce kidney injury, indicating by similar levels of plasmal cystatin C (1594.7+/- 88.74 ng/mL) compared with those under water feeding (1526.6+/- 109.28 ng/mL) (p=0.636)(Figure.9a).

**Figure 9.**
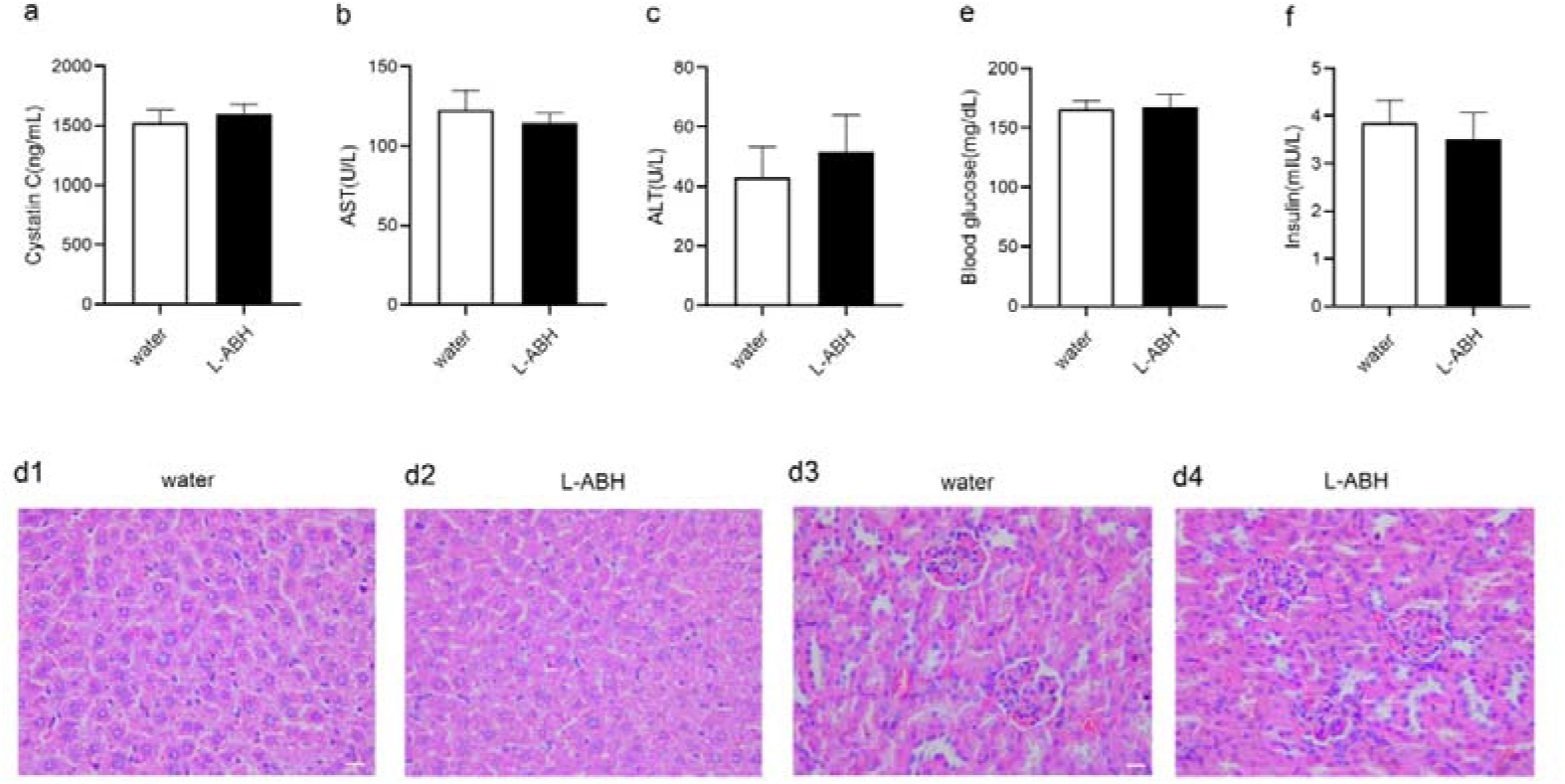
The effect of L-ABH on the liver and the kidney. WT C57BL/6 mice at the age of two months were subject to oral gavage of water(n=8) or L-ABH(n=8) for 12 weeks. The blood samples were collected for analyzing the levels of cystatin C(p=0.636)(**a**), AST(p=0.557)(**b**), and ALT(p=0.528)(**c**), relative to water feeding. The isolated liver and the kidney from water or L-ABH feeding were subject to H&E analyses. The images of the liver under water (**d1,**n=5) or L-ABH feeding(**d2,**n=5). The image of the kidney under water (**d3**,n=5) or L-ABH feeding (**d4**,n=5). Bar, 200 μm. Mice fasted O/N and blood samples were collected for measuring blood glucose(p=0.890)(**e**) or insulin levels (p=0.660)(**f**) from mice under water (n=8) or L-ABH feeding(n=8). Student’s *t*-test for differences (**a,b,c,e,f**).

L-ABH did not alter liver function, indicating by similar levels of plasmal aspartate aminotransferase (114.5+/-6.26 U/L) compared with those under water feeding(122.9+/- 12.45 U/L)(p=0.557) (Figure.9b). Similarly, L-ABH did not alter plasmal level of alanine aminotransferase(51.7+/- 12.31 U/L) compared with those under water feeding (43+/-10.26 U/L) (p=0.528) (Figure.9c). The isolated liver and kideny were processed into H& E staining. The microanatomy structures of the liver and the kidney were normal, indicating by intact liver lobule and renal corpuscle (Figures.9d1-9d4). Notably, L-ABH did not alter fasting blood glucose level of WT C57BL/6 male mice (167.6+/-10.71 mg/dL) compared with those under water feeding(165.8+/-7.05 mg/dL)(p=0.890)(Figure.9e). L-ABH did not alter fasting blood insulin level in WT C57BL/6 male mice (3.5+/-0.56 mIU/L) compared with those under water feeding(3.8+/-0.48 mIU/L)(p=0.660) (Figure.9f).

## Discussion

Laser photocoagulation and intravitreal injection of anti-VEGF antibodies or steroids are used to treat DR. These treatments alleviate symptoms by targeting retinal microvascular abnormalities but ignore the early pathologies, retinal neurodegeneration. Retinal neurodegeneration promotes DR progression from two aspects. On one hand, injured neurons release semaphorin 3A or erythropoietin, instigating pathological vascular permeability in diabetic retina [43, 44] . On the other hand, neuronal injury induces proliferation of Müller glia which release pro-inflammatory factor and VEGF, disrupting blood-retinal barrier [45, 46]. Thus, blockade of retinal neurodegeneration may stall DR at its very early stage.

In this study, we demonstrated that inhibition of SRR prevented DR through the neuroprotection in the retina as well as through maintaining glucose homeostasis in the whole body. Excitotoxicity is largely dependent on NMDAR which necessitates the presence of glutamate and its co-agonists, D-Ser or glycine [47, 48]. Diabetes impairs glutamate transport via glia [24, 25] but we only detected trace level of glutamate in the aqueous humor of db/db and of BKS WT mice. This inconsistency may result from fast glutamate metabolism via glutmate-glutamine-GABA cycle in the mice with BKS background. Intravitreal injection of L-ABH significantly reduced glutamate-induced neurotoxicity which significantly contributes to retinal neurodegeneration in DR [24–26]. The protection in the retina suggests that L-ABH decreases D-Ser levels in the retina. However, oral gavage of L-ABH did not reduce D-Ser levels in the aqueous humor of db/db mice, relative to db/db mice fed with water (Supplemental Figure 6). Loss-of-function mutation of *srr* only reduces D-Ser levels in the aqueous humor to ∼70% of those in WT mice [31], suggesting that there are sources synthesizing D-Ser other than SRR in the retina [49, 50].

We have shown that *srr* deletion and L-ABH treatment reduces production of pro-inflammatory factors and VEGF in retinal pigment epithelial cells [34, 51]. In our unpublished results, we demonstrated that deletion of *srr* in Müller cell reduced the transcription of *il-1*β and *il-18* (upon request). Thus, oral gavage of L-ABH presumably reduced production of inflammatory factors in the retina. We project that inhibition of SRR by L-ABH not only reduces D-Ser but also pro-inflammatory factors in the retina, which synergistically protect diabetic retina.

With db/db mice and DIO mice, application of L-ABH improved glucose tolerance and insulin sensitivity whereas L-ABH failed to normalize glucose homeostasis in streptozotocin-injected DIO mice(data not shown). These data suggest that SRR is a new target for modulating glucose homeostasis and insulin sensitivity in T2DM. At least, we have demonstrated that L-ABH normalized blood glucose through inhibiting liver gluconeogenesis. Comparing the difference of SRR expression between the liver and pancreatic islets, we considered that inhibition of SRR regulated glucose homeostasis mostly via inhibiting liver gluconeogenesis, thereby reducing hepatic glucose output in db/db mice. Inhibition of SRR by use of L-ABH did not alter GSIS in isolated pancreatic islets. Consistent with which, L-ABH did not alter serum insulin level in db/db mice, relative to water feeding. By contrast, deletion of *srr* increases GSIS *in vivo* [41]. Previous study indicate that NMDA-R antagonist, MK-801, increases GSIS in pancreatic β cells by prolonging depolarization and plateau calcium oscillations [52]. By inhibiting D-serine synthesis, L-ABH presumably blocks NMDAR in part in pancreatic β cells which is in contrast with complete blockade of NMDAR by MK-801. The difference possibly explains that L-ABH failed to modify GSIS. Hyperglycemia modifies amine group of proteins by reactive dicarbonyls, leading to formation of advanced glycation end products (AGEs) which are toxic to retinal neurovasaculature [53, 54] . Thus, normalization of glucose level in db/db mice by L-ABH may reduce the amount of AGEs which were not yet examined.

In summary, we indicate that inhibition of SRR is a novel target to manage DR and the effects of SRR inhibition on DR was mediated by the local and systemic effects.

## Acknowledgments

The study is supported by Integrated Project of State Key Laboratory of School of Optometry and Ophthalmology(#J02-20190204), Wenzhou Medical University, Wenzhou Municipal Scientific and Technology Bureau(#H20210012 and #Y20190062), National Natural Science Foundation of China for Youth (#82301614 and # 32000437), and Regional Innovation and Development Joint Fund of National Natural Science Foundation of China (# U23A20164).

The authors sincerely appreciate patch-clamp study from Prof. Chaoran Ren and Dr. Xiaodan Huang from Jinan University, Guangzhou.

## Abbreviations

ERG, electroretinogram; SD-OCT, spectral-domain optical coherence tomography; T2DM, type II diabetes; L-ABH, l-aspartic acid β-hydroxymate; SRR, serine racemase; G6Pase, glucose- 6-phosphatase; PEPCK, phosphoenolpyruvate carboxykinase; GSIS, glucose stimulated insulin secretion; DIO, diet-induced obesity; NMDA, N-methyl-D-aspartate; GCGR, glucagon receptor; OGTT, oral glucose tolerance test; ITT, insulin tolerance test

## Authors’ contribution

HY Jiang, PS Zhou, X Jiang conducted all the essential experiments, collected and analyzed the data; X Wu and YD Mao helped animal husbandry; SY Liu and SQ Tang helped HPLC; ZW Zhang and J Zhou examined liver and muscle glycogen; B Ren and G Shan provided reagents; J Qu provided resources; SZ Wu conceived of the project,analyzed the data,supervised the project,wrote the manuscript. SZ Wu takes the responsibility for the contents of the article.

