## Supplemental information for "Inhibition of serine racemase prevents diabetic retinopathy"

Correspondence should be directed to:

Prof. Shengzhou Wu, Ph.D, M.D

Orcid# [0000-0003-1154-2369](https://orcid.org/0000-0003-1154-2369)

Fax：(86)-577-88067934

**Supplemental methods**

**Comprehensive lab animal monitoring system (CLAMS)**

CLAMS cages(Oxymax; Columbus Instruments, Columbus, OH) were used to monitor metabolism including consumption of food, water, and oxygen, production of carbon dioxide, respiratory quotient, and sleep episodes. The db/db mice feeding with water, L-ABH, and metformin and BKS WT mice were evaluated with CLAMS cages at the end of feeding. Mice were housed individually in animal facility of School of Optometry and Ophthalmology in Wenzhou Medical University with water and food ad libitum under a 12/12-h light/dark cycle. The mice were acclimated for one day before data collection, and data were collected for three consecutive days.

**Glucose stimulated insulin secretion (GSIS) assay**

Pancreatic islets from adult WT C57B/6 mice were isolated and cultured in 24-well plate. The islets were starved for 1h in Krebs-Ringer bicarbonate HEPES buffer or in the solution added with L-ABH(125 μM) and were switched into Krebs-Ringer bicarbonate HEPES buffer added with 2 mM D-glucose. Following the operation, the cultures were subject to sham treatment or L-ABH for 1 h(125 μM). The supernatants were collected for measuring insulin contents. To measure insulin secretion under high concentration of D-glucose, the media were switched into Krebs-Ringer bicarbonate HEPES buffer added with 20 mM D-glucose and treated for 1 h with vehicle or L-ABH(125 μM). The supernatants were collected for measuring insulin contents. The secreted insulin was expressed as the ratios between insulin in the supernatants and total protein mass in the lysates.

**Supplemental figures and figure legends**

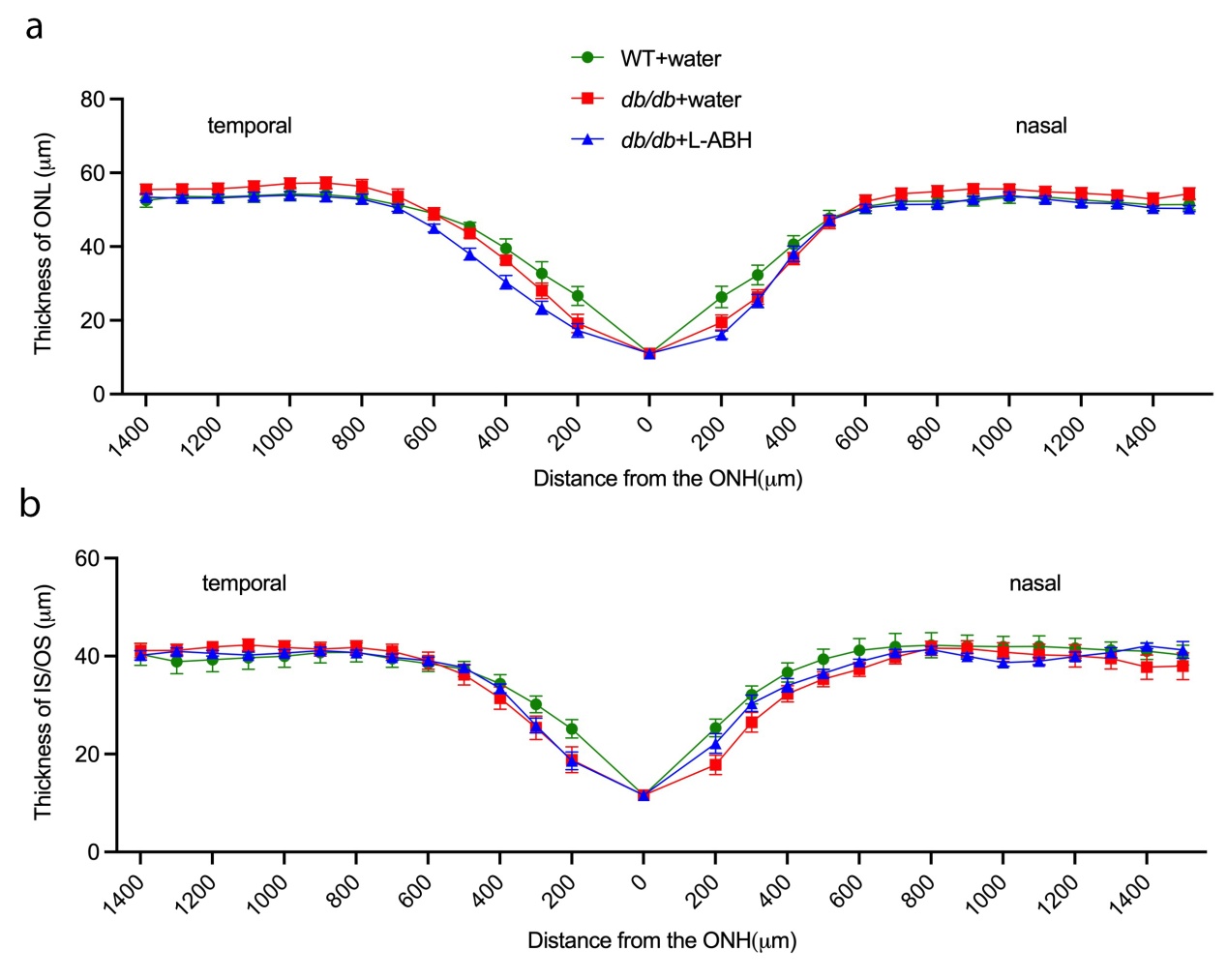

**Supplemental Figure 1. L-ABH did not alter the thickness of outer retina.**

Db/db male mice were fed for 18 weeks with water (db/db+water) or L-ABH (20 mg/kg/d) (db/db+L-ABH). BKS WT mice on regular chow diet were included (WT). At the end of feeding, the mice were subjected to SD-OCT examination. Under linear mode of scan, the optic disk was positioned in the center of the scan box and the images were automatically acquired from the nasal to the temporal retina. The thickness of ONL (**a**) and IS/OS (**b**) were recorded under 200 μm interval with reference to optic nerve head (ONH) as the central point. The quantifications were averaged from 6-10 mice.

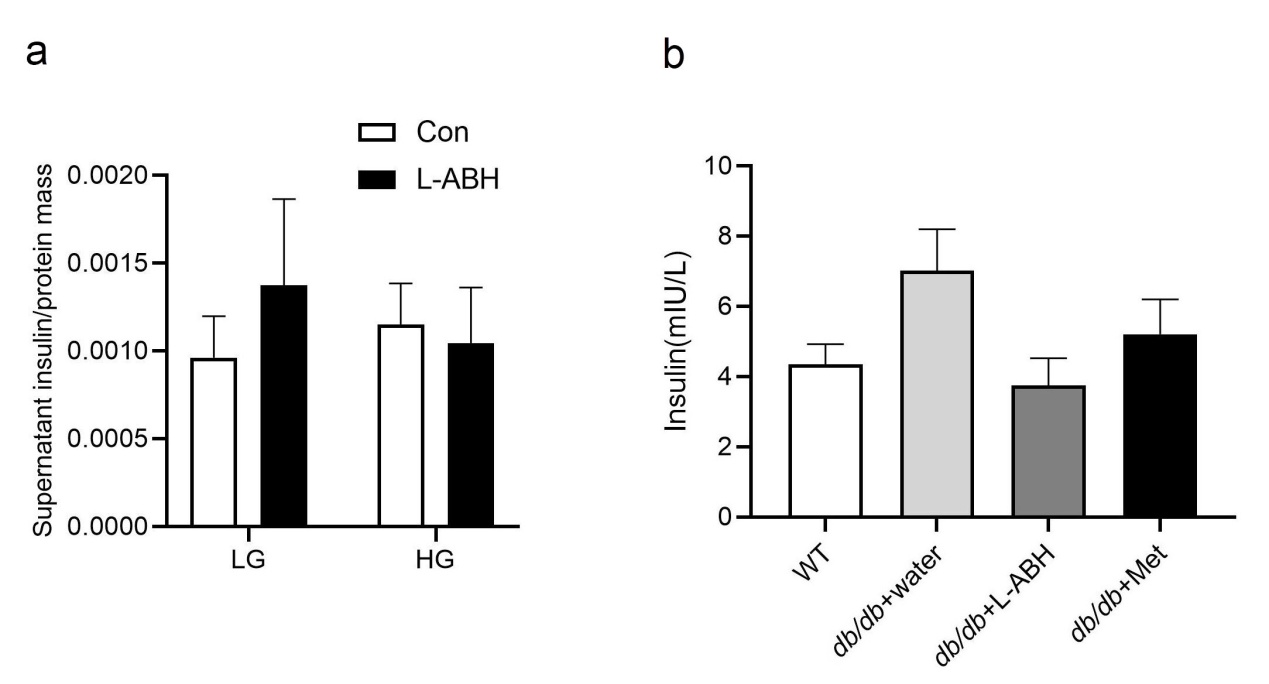

**Supplemental Figure 2. L-ABH did not alter GSIS *in vivo and in vitro*.** (**a**) Insulin secretion examination in islet cultures. LG indicated 2 mM glucose and HG indicated 20 mM glucose in culture media. (**b**) The mice fasted O/N and retroorbital blood collection was conducted at the end of feeding for WT(n=14), db/db +water(n=15), db/db+ L-ABH(n=10), and db/db+metformin (n=9) and subject to insulin examination by ELISA. One-way ANOVA with repeated measures for inter-treatment evaluations.

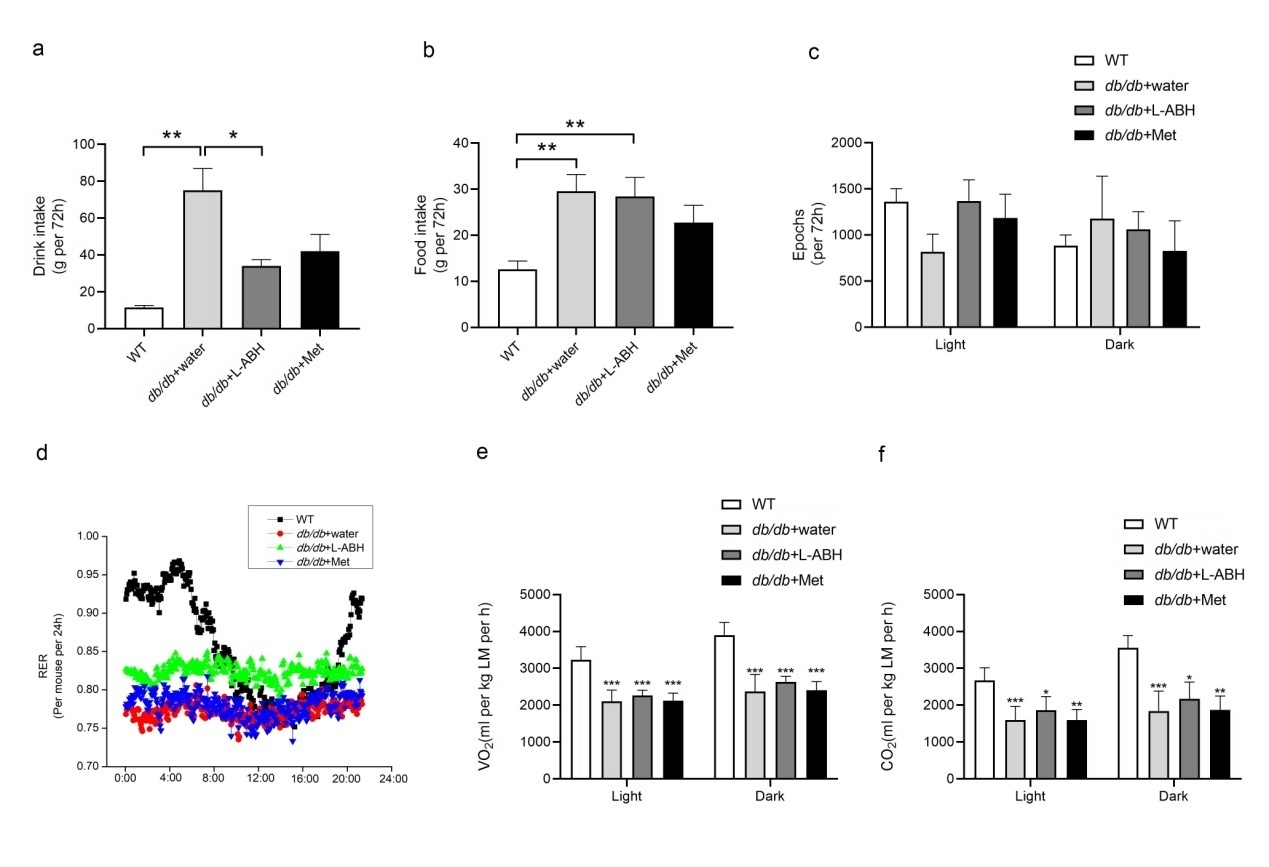

**Supplemental Figure 3. CLAMS cage analysis.** At 18 weeks of feeding, BKS WT mice(n=10) and db/db mice under water feeding(n=10), L-ABH feeding(n=10),metformin feeding(n=6) were subject to CLAMS assay. (**a**)Water consumption. (**b**) Food consumption. (**c**) Locomotor activity. (**d**) Respiratory quotient. (**e**)Oxygen consumption. ***p<0.001 indicated differences between feeding of either chemical and water, respectively (**f**)Production of carbon dioxide. *p<0.05,**p<0.01, and ***p<0.001 indicated differences between feeding of either chemical and water, respectively. One-way ANOVA with repeated measure for differences (**a,e,f**-dark). Kruskall-Wallis for differences(**b,f**-light). *p<0.05, **p<0.01 indicated differences between indicated groups.

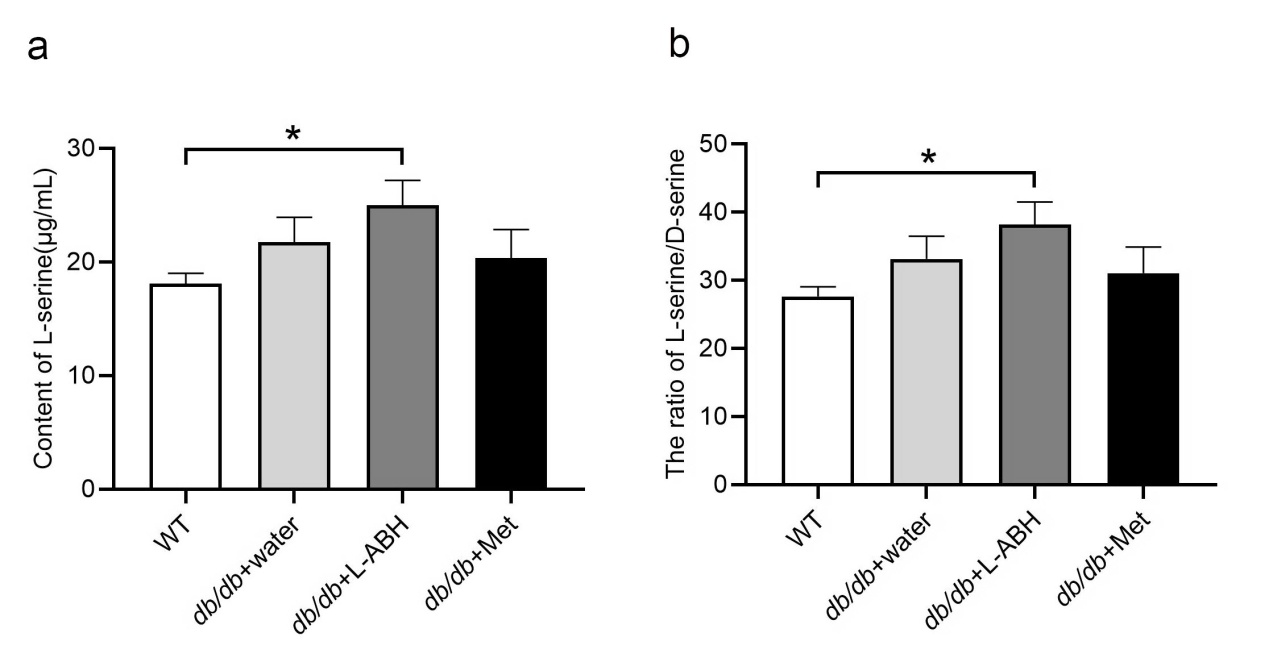

**Supplemental Figure 4. rp-HPLC analysis of serum l-/D-serine. (a)** At the end of feeding, blood sample were collected from BKS WT mice(n=6) and db/db mice feeding with water(n=7),L-ABH(n=8),metformin(n=6) and the samples were subject to rp-HPLC analysis. (**b**) Quantification of l-/D-serine ratios in **a**. One-way ANOVA with repeated measures for differences. *p<0.05 indicated differences between indicated groups.

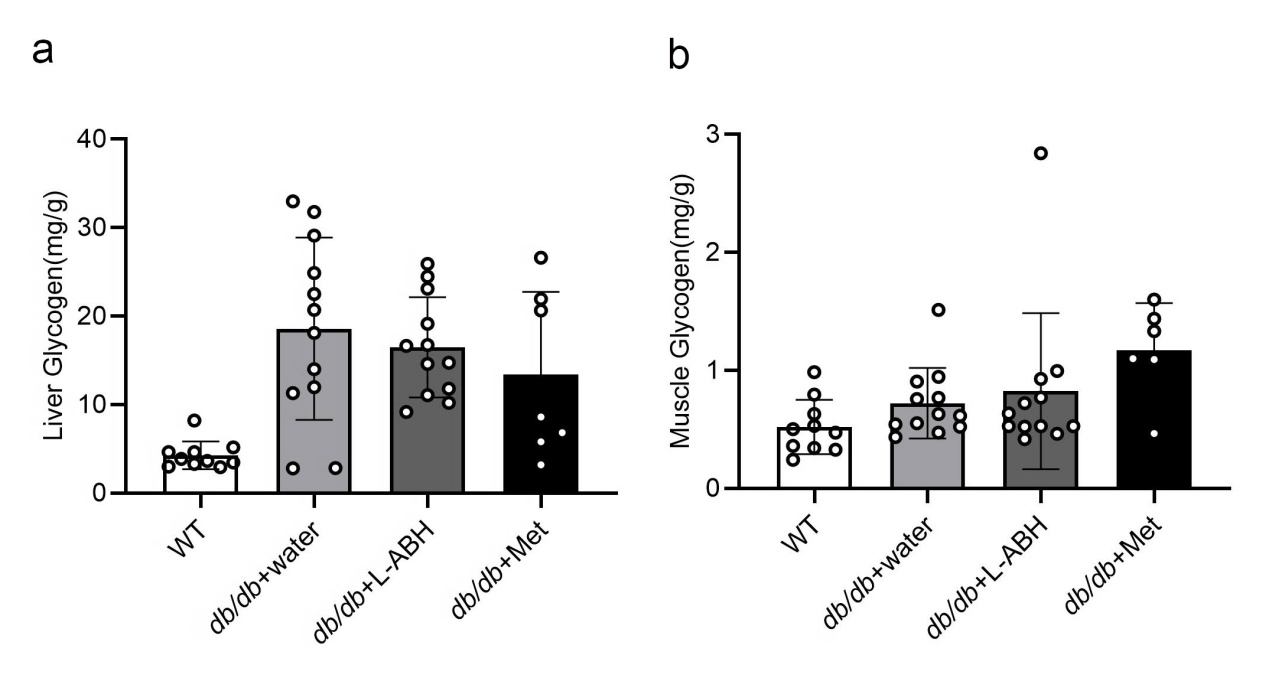

**Supplemental Figure 5. Glycogen levels in the liver and in the gastrocnemius muscle in db/db mice feeding with water and L-ABH.** At the end of feeding,the liver and gastrocnemius muscle were isolated from BKS WT mice(n=10) and db/db mice under the feeding of water(n=13), L-ABH (n=12),metformin(n=6). The liver (**a**) and muscle tissues (**b**) were homogenized and subjected to glycogen analysis with a commercial kit (Solarbio,Beijing). One-way ANOVA with repeated measures for differences.

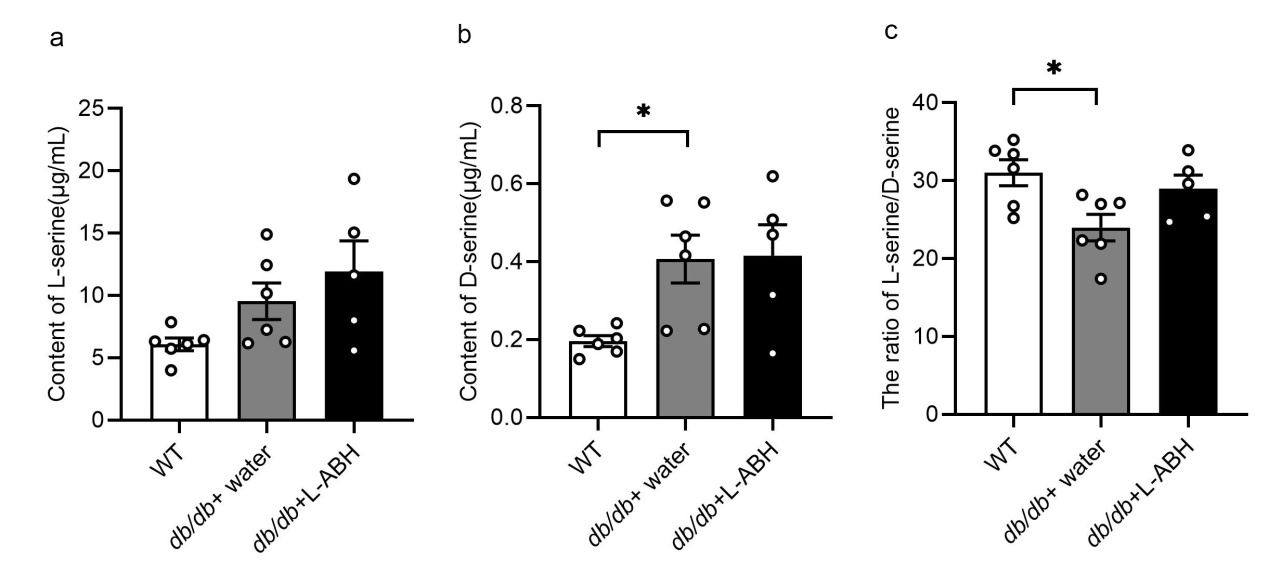

**Supplemental Figure 6. rp-HPLC analysis of aqueous humor l-/D-serine.** At the end of feeding, the aqueous humor were collected from BKS WT mice(n=6) and db/db mice feeding with water(n=6),L-ABH(n=5) and the samples were subject to l-serine (**a** ) and D-serine (**b**) analyses by rp-HPLC. (**c**) Quantification of l-/D-serine ratios. One-way ANOVA with repeated measures for differences. *p<0.05 indicated differences between indicated groups.
